# Distinct roles of three trypanosomal Oxa1 insertases in biogenesis of mitochondrial membrane complexes

**DOI:** 10.64898/2026.06.30.735475

**Authors:** Jonathan Espinoza Wong, Ingrid Škodová-Sveráková, Jan Říha, Prashant Chauhan, Alexandra List, Verena Danzinger, Alena Zíková, Ondřej Gahura

## Abstract

The insertase Oxa1 is required for protein insertion into the inner mitochondrial membrane and for the biogenesis of oxidative phosphorylation complexes. While most eukaryotes encode one or two Oxa1 proteins, we identified three paralogs in *Trypanosoma brucei*: TbOxa1-1, TbOxa1-2, and TbOxa1-3. Knock-out of individual paralogs followed by phenotypic analyses and proteomic characterization of submitochondrial fractions revealed distinct functions. Respiratory chain complexes I and IV are primarily affected by loss of TbOxa1-1, whereas complex III and ATP synthase depend on TbOxa1-2; the ablation of TbOxa1-3 results in minor phenotypes in culture. In TbOxa1-2-depleted cells, ATP synthase biogenesis is compromised by the defective import or processing of the nuclear-encoded subunit-c, which also requires a rhomboid peptidase-like protein. Further, the ablation of TbOxa1-2 triggers accumulation of membrane proteins in the matrix, supporting its role in conservative sorting. Together, our results demonstrate that the trypanosomal Oxa1 machinery evolved a paralog-specific division of labor to manage a highly divergent mitochondrial membrane proteome.

## INTRODUCTION

Translocation of proteins across membranes, their insertion into membranes and biogenesis of membrane complexes in eukaryotic cells involve conserved machineries, including components of bacterial origin. Proteins encoded in the genomes of mitochondria, endosymbiotic organelles, are inserted into the lipid phase of the inner mitochondrial membrane by a protein first reported in *Saccharomyces cerevisiae* and termed <u>ox</u>idase <u>a</u>ssembly 1 (Oxa1^1^). The mitochondrial insertase Oxa1 and its chloroplastic counterpart Alb3 are homologous to bacterial YidC, which acts in the Sec-translocon dependent and independent protein targeting into the membrane^2,3^. In addition, three divergent Oxa1 homologs have been identified and structurally characterized as components of three distinct complexes for protein transport to endoplasmic reticulum in yeast and humans^4^, documenting that the role of the proteins from Oxa1-superfamily is not limited to bacteria and endosymbiotic organelles.

Although mitochondrial Oxa1 is ubiquitous in eukaryotes^5^, so far it has been studied predominantly in *S. cerevisiae* and human cells (^6^ and references below; reviewed in ^7^ and ^8^). Budding yeast Oxa1 facilitates co-translational insertion of mitochondrially encoded nascent polypeptides and post-translation assembly of oxidative phosphorylation (OXPHOS) complexes, i.e. electron transport chain complexes I to IV (cI to cIV) and ATP synthase, that contain mitochondrially encoded subunits^9–11^, consistently with its bacterial origin. Oxa1-facilitated co-translational protein folding and insertion was recently visualized in human cells^12^. Oxa1 in *S. cerevisiae* is also required for incorporation of several nuclear encoded proteins into the inner mitochondrial membrane by the conservative sorting pathway^13,14^. Substrates of this pathway, exemplified by a precursor of Oxa1 itself, ABC transporters, or TIM22 carrier translocase subunits Tim18 and Sdh4^14–17^, contain transmembrane segments that are first imported into the mitochondrial matrix and re-inserted into the membrane by Oxa1 from inside. However, whether this Oxa1 function is specific to yeast or more widespread in eukaryotes remains unknown. Eukaryotic genomes typically encode a single Oxa1 protein, but some groups have two Oxa1 paralogs. In addition, most groups have a divergent paralog, referred to as Oxa2 in humans and plants and Cox18 in *S. cerevisiae*, which most likely branched from Oxa1 before the last eukaryotic common ancestor^5,18^. Oxa2/Cox18 is involved in the biogenesis of cIV in animals, fungi and plants by facilitating the insertion of the Cox2 subunit^18–21^.

The parasitic protist *Trypanosoma brucei* belongs to the class Kinetoplastida within an early-branching eukaryotic supergroup Discoba and features numerous non-canonical cellular processes and structures^22^. Among others, the trypanosomal inner mitochondrial membrane complexes, including the OXPHOS system and protein import complexes, are markedly divergent from those of classical model organisms^23–26^, implying differences in their assembly. The parasite’s mitochondrial encoded proteins are highly hydrophobic, partially due to extensive uridine insertion/deletion editing of their primary transcripts ^27^, which results in exceptionally phenylalanine- and leucine-rich sequences. Therefore, they presumably require adaptations of the machinery for their insertion. In budding yeast, several factors cooperate with Oxa1 in the cotranslational insertion of mitochondrial encoded proteins to the membrane^11,28^. So far, only a homolog of one of these factors, Mba1, has been identified in *T. brucei*. Mba1 in yeast acts as a mitochondrial ribosome receptor, which facilitates the attachment of the translating mitoribosomes to the membrane via Oxa1^29,30^. However, although it appears to play a role in the biogenesis of OXPHOS complexes in *T. brucei*, it is not essential for the tethering of mitoribosomes to the membrane^31^. Furthermore, a large proportion of nuclear encoded inner mitochondrial membrane proteins in *T. brucei* appear to be delivered to the membrane via the conservative pathway^32^. Therefore, mechanisms and players of the insertion of proteins into the inner mitochondrial membrane phospholipid bilayer and assembly of membrane complexes in *T. brucei* presumably differ from model eukaryotes.

The trypanosomal toolkit for membrane biogenesis is assumed to involve the insertase Oxa1, the presence of which was inferred based on sequence similarity^5,31^ but has never been addressed experimentally. Here, we show that the inner mitochondrial membrane in *T. brucei* contains three divergent homologs of Oxa1, which are differentially involved in biogenesis of OXPHOS complexes with mitochondrial encoded subunits. A comprehensive analysis of submitochondrial proteomes further reveals a role of one of the three trypanosomal Oxa1 paralogs in the conservative sorting of nuclear-encoded membrane proteins and in ATP synthase subunit c processing, which is facilitated by a rhomboid protease-like cofactor.

## RESULTS

### The genome of *Trypanosoma brucei* encodes three paralogs of Oxa1

To identify candidates for Oxa1 in *Trypanosoma brucei*, we searched its predicted proteome by protein BLAST with human and *S. cerevisiae* Oxa1 sequences as queries and found two Oxa1-like proteins. Using the sequence of one of them as a query, we identified a third candidate. The same three candidates were detected by structure similarity search in the AlphaFold database by FoldSeek queried by *Bacillus halodurans* YidC structure resolved by X-ray crystallography (PDB ID 3WO7^33^). Thus, by sequence- and structure- similarity based approaches, we identified three putative mitochondrial Oxa1 insertases in the nuclear genome of *Trypanosoma brucei*, hereinafter referred to as TbOxa1-1 (TriTrypDB gene ID Tb927.11.6150), TbOxa1-2 (Tb927.9.10050), and TbOxa1-3 (Tb927.5.2030).

Although the three paralogs exhibit low sequence similarity when compared to each other, and to mammalian and yeast Oxa1, and bacterial YidC (Supplementary Fig. 1), the Alphafold predicted structures indicate that the candidates are Oxa1 homologs. All of them contain a core hydrophobic domain of five transmembrane helices typical for the Oxa1 family flanked by soluble N- and C-terminal segments, exposed presumably to intermembrane space and mitochondrial matrix, respectively (Fig. 1A). The positively charged arginine residue found in the hydrophilic groove between the transmembrane helices of the bacterial YidC (R72 in *Bacillus subtilis* YidC) and implicated to be crucial for its membrane insertion activity^33,34^ is conserved in all three trypanosomal paralogs. At least one of two proximally situated aromatic residues contributing to substrate binding (Y243 and W244 in YidC) are also present, supporting a role in membrane protein insertion (Fig. 1B,C & Supplementary Fig. 1). In yeast and mammals, the C-terminal tail of Oxa1 is exposed to matrix and binds translating mitochondrial ribosomes in the vicinity of the exit from their polypeptide tunnel^35,36^. The length and sequence of the C-terminal segments vary between the three TbOxa1 proteins, suggesting that if they interact with mitoribosomes, the nature or extent of these interactions may differ among them. The C-terminal tail of TbOxa1-1 contains a region that is similar to the mitoribosome binding site 2 of human Oxa1^36^ (Supplementary Fig. 1). The extended and divergent C-termini of TbOxa1-2 and TbOxa1-3 might imply substrate specificity or different interactions with protein import or OXPHOS assembly toolkits.

**Figure 1.**
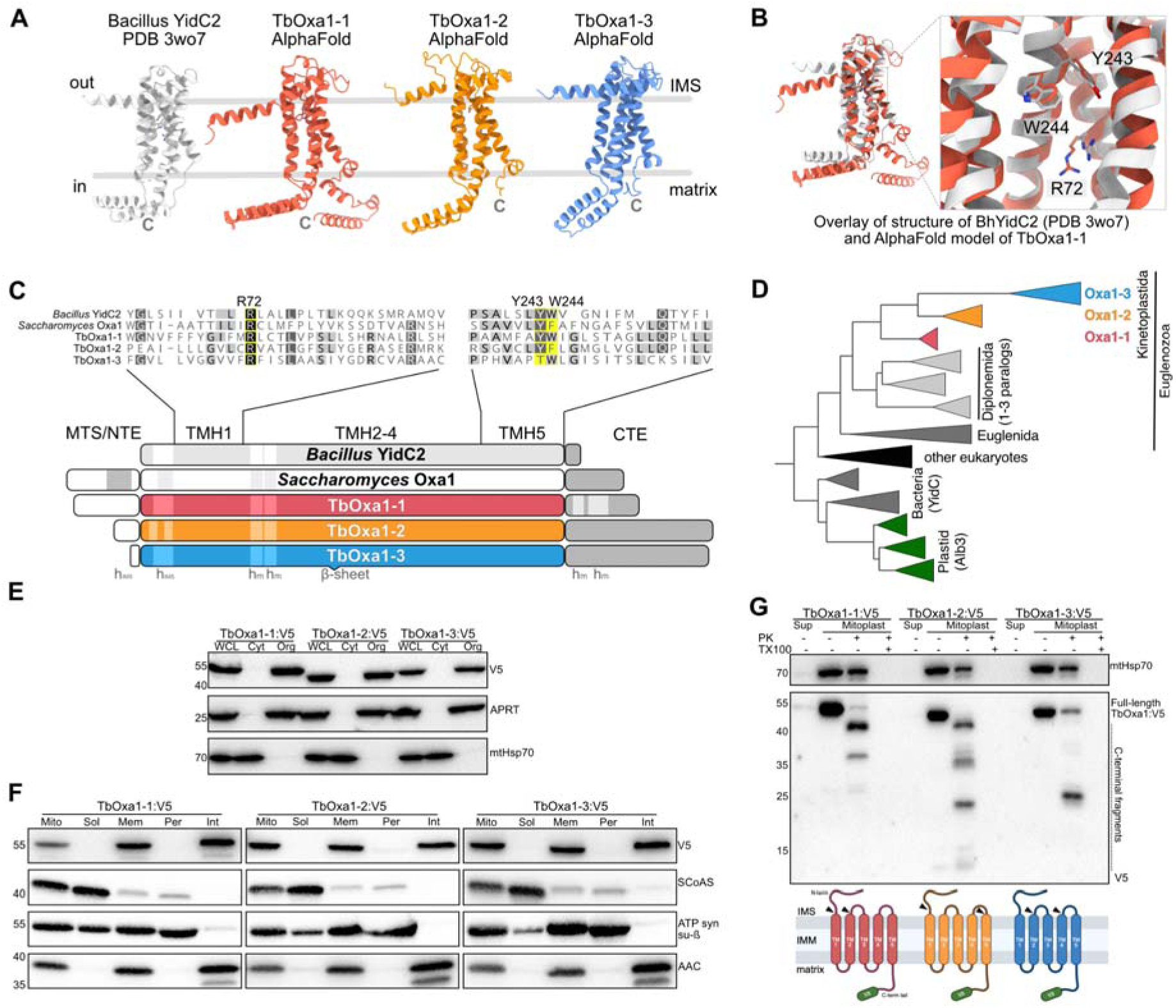
*T. brucei* possesses three Oxa1 paralogs with canonical topology in the inne mitochondrial membrane. **(A)** AlphaFold models of the TbOxa1 proteins and the crystal structure of *Bacillus halodurans* YidC2 (PDB 3WO7). IMS - intermembrane space. **(B)** Superposition of AlphaFold model of TbOxa1-1 (red) with structure of BhYidC2 (white) with conserved residues crucial for BhYidC2 insertase activity shown as sticks. Numbering is based on BhYidC2. **(C)** Schematic representation of TbOxa1 proteins, *Saccharomyces cerevisiae* Oxa1 (ScOxa1), and BhYidC2. TMH - transmembrane helix; h_m_ and h_IMS_ - matrix- and IMS-exposed helices; MTS - mitochondrial targeting sequence; NTE and CTE N- and C-terminal extensions. Alignment of sequences of TMH1 and TMH5 is shown with the residues depicted in panel (B) highlighted in yellow. **(D)** A collapsed phylogenetic tree of Oxa1 homologs showing two duplication events in Kinetoplastida. **(E)** Immunoblots of lysates from whole cells (WCL) and cytosolic (Cyt) and organellar (Org) fractions generated from V5-tagged TbOxa1-expressing *T. brucei* (TbOxa1:V5) cells. **(F)** Immunoblots of lysates from mitochondria (Mito) isolated from TbOxa1:V5 cells and respective submitochondrial fractions (Sol - soluble, Mem - total membrane, Per - peripheral membrane, Int - integral membrane). **(G)** Immunoblots of lysates from isolated mitoplasts differentially treated with proteinase K (PK) and Triton-X (TX100). A scheme shows the sites cleaved by proteinase K (arrowheads) resulting in V5-bearing C-terminal fragments. Sup - supernatant after mitoplasts isolation. For panels (E) to (G), we used primary antibodies to detect V5-tagged TbOxa1 proteins and marker proteins of individual fractions: APRT (Cyt), Hsp70 (Mito), SCoAS (Sol), ATP synthase subunit β (Per), and ADP/ATP carrier (AAC; Int).

Eukaryotes typically possess one, in some cases two, Oxa1 proteins. Thus, the occurrence of three Oxa1 paralogs in kinetoplastids, previously suggested^5^, is unusual. To shed light on the evolutionary history of the three Oxa1 genes in *T. brucei*, we searched predicted proteomes of selected euglenozoans for Oxa1 and reconstructed phylogenetic tree with the obtained sequences and Oxa1-superfamily representatives from other eukaryotes and bacteria (Supplementary Fig. 2). The analysis revealed that all three Oxa1 paralogs are found only in trypanosomatids and the sister lineage bodonids (together Metakinetoplastina), while two are present in representatives of the early branching kinetoplastid group Prokinetoplastina. Thus, the first and second Oxa1 gene duplication, the former separating TbOxa1-1 and the latter resulting in TbOxa1-2 and TbOxa1-3, occurred in the common ancestor of Kinetoplastida and Metakinetoplastina, respectively (Fig. 1D & Supplementary Fig. 2; see legend of Supplementary Fig. 2 for details on the results of the analysis). Therefore, any possible functional implication associated with the three paralogs of Oxa1 is restricted to trypanosomatid parasites and closely related species.

### TbOxa1 proteins are embedded in the inner mitochondrial membrane with C-termini exposed to matrix

To determine subcellular localization and membrane topology of TbOxa1 proteins, we modified one allele of each TbOxa1 by homologous recombination in the procyclic *T. brucei* to generate strains expressing individual TbOxa1 proteins with a V5-epitope at their C-termini. The introduced V5 epitopes did not interfere with cell viability (Supplementary Fig. 3A). Solubilization of cells by the mild detergent digitonin followed by differential centrifugation showed that all TbOxa1 homologs are present exclusively in the organellar fraction, indicating mitochondrial residency (Fig. 1E). Further, fractionation of isolated mitochondria by differential centrifugation and alkaline carbonate treatment documented that the TbOxa1 proteins integrate into the inner mitochondrial membrane (Fig. 1F).

To address the orientation of TbOxa1 proteins in the membrane, mitoplasts isolated from *T. brucei* cells harboring V5-tagged TbOxa1 proteins were subjected to proteinase K protection assay ^37^. The results showed that the V5-tagged C-terminal tails were protected from proteolytic degradation in intact mitoplasts, consistent with the canonical Oxa1 N-out/C-in orientation (Fig. 1G). The electrophoretic mobility of the individual TbOxa1 proteins under native conditions indicated that all paralogs are found at least partially in complexes of molecular weight estimated to approximately 480 kDa (Supplementary Fig. 3B), indicating association with other protein factors.

Together, these results document that all three trypanosomal Oxa1 have canonical localization and topology in the inner mitochondrial membrane, consistent with their presumed role in the membrane biogenesis.

### TbOxa1-2 is required for the ATP synthase biogenesis in bloodstream form trypanosomes

In model eukaryotes, the primary function of Oxa1 is the insertion of mitochondrially encoded subunits of OXPHOS complexes in the inner mitochondrial membrane^9–11^. The life cycle stage of *T. brucei* residing in the blood of mammalian hosts lacks functional cI, cIII and cIV of the electron transport chain and rely on the reverse proton pumping activity of ATP synthase for the maintenance of the mitochondrial membrane potential^38–40^. Consequently, subunit-a of ATP synthase is the only essential mitochondrial encoded membrane protein, rendering bloodstream *T. brucei* a unique model to study the role of the TbOxa1 paralogs.

Using the CRISPR/Cas9, we knocked-out both alleles of individual TbOxa1 genes (Supplementary Fig. 4A,B). All three bloodstream form knock-out strains exhibited no (doubling time 7.4 hours in both the TbOxa1-2^KO^ cells and the parental strain) to mild (8.5 hours in TbOxa1-1^KO^ and 9.0 hours in TbOxa1-3^KO^ cells) growth defects (Fig. 2A). However, native electrophoresis of mitochondrial lysates followed by immunoblotting with antibodies against ATP synthase subunits revealed a decreased abundance of ATP synthase complexes, most prominent in the TbOxa1-2^KO^ strain (Fig.2B). Remarkably, the absence of TbOxa1-2 resulted in a complete loss of a band, which represents a subcomplex of the catalytic matrix-exposed part, F_1_-ATPase, associated with the c_10_-ring (F_1_+c_10_-ring^25^). Subunit-c is a known substrate of YidC in bacteria and Oxa1 in *S. cerevisiae*^41,42^. Unlike the yeast subunit-c, which is encoded in the mitochondrial genome, the gene of the trypanosomal counterpart has been transferred to the nuclear genome. The loss of F_1_+c_10_-ring band suggests that c_10_-ring assembly or its association with the rest of the complex depends on TbOxa1-2 despite subunit-c being imported from the cytosol. The reduced abundance of ATP synthase was further confirmed by measurement of its oligomycin-sensitive proton pumping capacity in isolated mitochondria by safranin-O assay. Consistently with the results of native immunoblots, the pumping activity of ATP synthase was affected in all strains, and most decreased upon deletion of TbOxa1-2 (Fig.2C).

**Figure 2.**
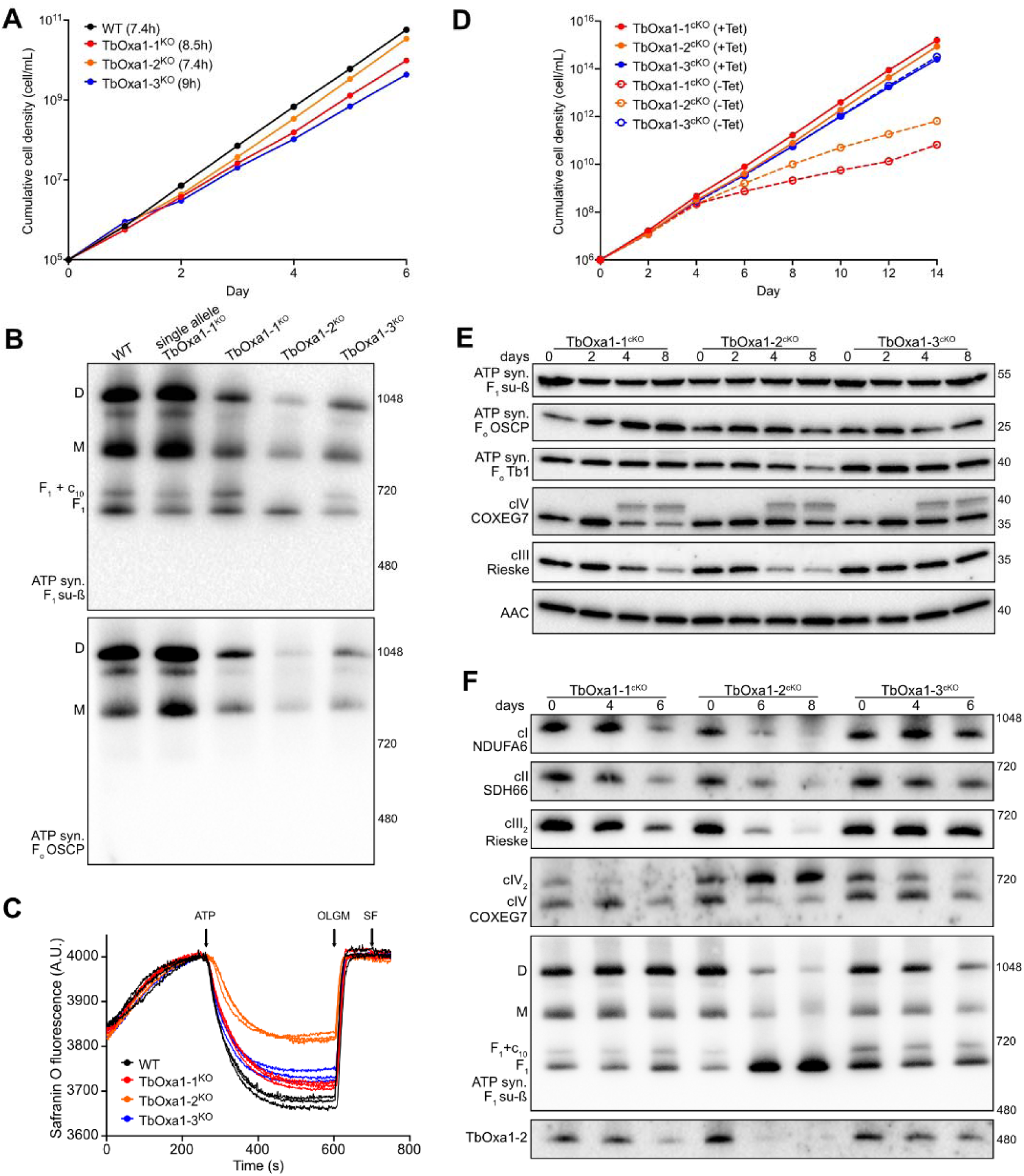
Knock-out of TbOxa1 paralogs differentially affects viability and OXPHOS complexes in bloodstream and procyclic trypanosomes. **(A)** Growth curves of Cas9-expressing parental (WT) and derived TbOxa1 knockout (KO) bloodstream form cell lines. **(B)** Immunoblots of mitochondrial lysates from WT and TbOxa1^cKO^ bloodstream form cells resolved by BN-PAGE probed with antibodies against ATP synthase subunits β and OSCP. **(C)** Mitochondrial membrane polarization capacity of digitonin-permeabilized WT and TbOxa1^cKO^ bloodstream form cells measured by safranine O staining. Arrows indicate addition of ATP, oligomycin (OLG), and protonophore SF6847 (SF). **(D)** Growth curves of TbOxa1^cKO^ procyclic form cell lines grown in SDM-79 medium with the ectopic allele switch on (+Tet) o off (-Tet). **(E)** Immunoblots of whole cell lysates from TbOxa1^cKO^ procyclic cells harvested on indicated days after tetracycline removal resolved by SDS-PAGE. **(F)** Immunoblots of mitochondrial lysates from TbOxa1^cKO^ procyclic cells harvested on indicated days after tetracycline removal resolved by BN-PAGE. For panels (E) and (F), we used primary antibodies to detect subunits of OXPHOS complexes and other mitochondrial proteins: TbOxa1-2, AAC, cI subunit NDUFA6, cII subunit SDH66, cIII subunit Rieske, cIV subunit COXEG7, and ATP synthase subunits β, OSCP and Tb1.

The rather weak growth phenotypes of individual TbOxa1 knock-outs might indicate functional redundancy among the paralogs. To test this hypothesis, we generated cell lines with all pairwise combinations of TbOxa1 paralogs knocked out. The cell line with knock-out of TbOxa1-1 and TbOxa1-3 exhibited a slower growth than individual knock-outs (doubling time 11.0 hours) but remained viable. The other two knock-out pairs resulted only in a slight, if any, additive growth defect (Supplementary Fig. 4C). The decrease of ATP synthase levels was not more pronounced than in the individual knockouts (Supplementary Fig. 4D). The absence of growth defects in TbOxa1-2^KO^, whether alone or in combination with a knock-out of another paralog, is consistent with the observation that the cultured bloodstream form trypanosomes tolerate a substantial reduction in ATP synthase levels; only a drop to less than 10 % of normal levels is associated with growth retardation^43^.

We conclude that TbOxa1-2, and to a minor extent also other two paralogs, are required for efficient assembly of ATP synthase. Based on the band patterns on the native immunoblots, we reason that the defect might be caused by an inefficient membrane insertion of nuclear encoded subunit-c in TbOxa1-2^KO^. Possibly, the decreased ATP synthase levels in the absence of TbOxa1-1 and TbOxa1-3 might be a consequence of inefficient insertion of the mitochondrial encoded subunit-a or secondary phenotypes.

### The loss of individual TbOxa1 proteins differentially affects the abundance of oxidative phosphorylation complexes

To address the role of TbOxa1 proteins in the assembly of all OXPHOS complexes, we used the procyclic insect form of *T. brucei*, which requires OXPHOS complexes for ATP production. All OXPHOS complexes except for cII contain at least one mitochondrial encoded membrane subunit and therefore require Oxa1 for their biogenesis in model species^9–11^. We generated cell lines with individual TbOxa1 paralogs knocked-out by CRISRP/Cas9 in the presence of an ectopic copy of the respective paralog expressed from a tetracycline-inducible promoter (conditional knock-out cell lines TbOxa1-1^cKO^, TbOxa1-2^cKO^ and TbOxa1-3^cKO^). We verified the successful deletion and replacement of the endogenous genes by diagnostic polymerase chain reaction (PCR; Supplementary Fig. 5A,B) and showed that in all three cell lines the mRNA expressed from the ectopic conditional allele was efficiently downregulated by day two after tetracycline removal (Supplementary Fig. 5C). In addition, an antibody we raised against TbOxa1-2 documents efficient downregulation of the protein in whole cell lysate and organellar fraction four days after tetracycline removal (Supplementary Fig. 5D). The repression of the ectopic TbOxa1 genes in cKO strains following tetracycline removal did not increase the mRNA levels of the other two paralogs. Furthermore, protein levels of TbOxa1-2 in TbOxa1-1^cKO^ and TbOxa1-3^cKO^ cells remained unaffected (Supplementary Fig. 5C,D), indicating that the ablation of one TbOxa1 paralog is not compensated by the upregulation of expression of the other paralogs.

Knock-out of TbOxa1-1 and TbOxa1-2, but not that of TbOxa1-3, caused a strong growth defect onsetting on day four after tetracycline removal (Fig. 2D). Such growth phenotypes would be consistent with a deficiency in the OXPHOS system. We used a panel of antibodies to assay the levels of OXPHOS complexes and their subunits in the absence of individual TbOxa1 paralogs (Fig. 2E). Probing of whole cell lysates resolved by denaturing electrophoresis documented accumulation of precursors of COXEG7 (previously referred to as trCOIV), a subunit of cIV^44–46^ upon knock-out of all TbOxa1 paralogs. This observation suggests that the absence of TbOxa1 proteins affect import or proteolytic processing of the protein, albeit perhaps indirectly via its inefficient assembly into cIV. Further, we observed decreased amounts of protein Rieske, a subunit of cIII, in TbOxa1-1^cKO^ and TbOxa1-2^cKO^ cells, more pronounced in the latter strain.

Detection of subunits of ATP synthase documented a decrease in subunits ATPTB1 and OSCP, located in the membrane and peripheral stalk^25^, respectively, but not in subunit-β of the soluble F_1_-ATPase subcomplex^47^, in TbOxa1-2^cKO^ cells. Immunoblotting of mitochondrial lysates resolved by electrophoresis under native conditions (Fig. 2F) confirmed the loss of F_1_+c_10_-ring, monomeric and dimeric ATP synthase, and accumulation of F_1_-ATPase upon TbOxa1-2 knockout, consistent with results in the bloodstream trypanosomes. The same cell line showed marked decrease in cIII, partially observed also in TbOxa1-1^cKO^ cells. The levels of cIV were to some extent affected in all cKO strains, but the decrease was by far most pronounced in TbOxa1-1 knockout, as also documented by in-gel activity staining (Supplementary Fig. 5F). The abundance of cI and cII appeared to be decreased to similar levels in both TbOxa1-1^cKO^ and TbOxa1-2^cKO^ cells. Overall, these results indicate that individual TbOxa1 paralogs have differential effects on the abundance of OXPHOS complexes, presumably due to defects in their biogenesis.

### Determination of submitochondrial *T. brucei* proteomes by label-free mass spectrometry

To investigate the influence of the individual TbOxa1 proteins on the overall mitoproteome landscape, we optimized and validated an approach for fractionation of mitochondria followed by label-free quantitative mass spectrometry (MS) to analyze the protein content of mitochondrial compartments. Isolated mitochondria were permeabilized and fractionated into soluble and total membrane fractions, and the content of the latter was subfractionated to peripheral and integral membrane fractions by sodium carbonate extraction (Fig. 3A; see Methods for details;^48^). The submitochondrial fractions were analyzed by denaturing electrophoresis and immunoblotting using antibodies against selected mitochondrial proteins with known function and localization. Specifically, we probed soluble matrix enzyme succinyl coenzyme A synthetase (SCoAS), a soluble intermembrane space protein cytochrome c, integral proteins of the outer and inner membrane, VDAC and ATP/ADP carrier (AAC), respectively, and several integral and peripheral components of OXPHOS complexes. All the marker proteins showed expected fractionation patterns, validating the approach (Fig. 3B).

**Figure 3.**
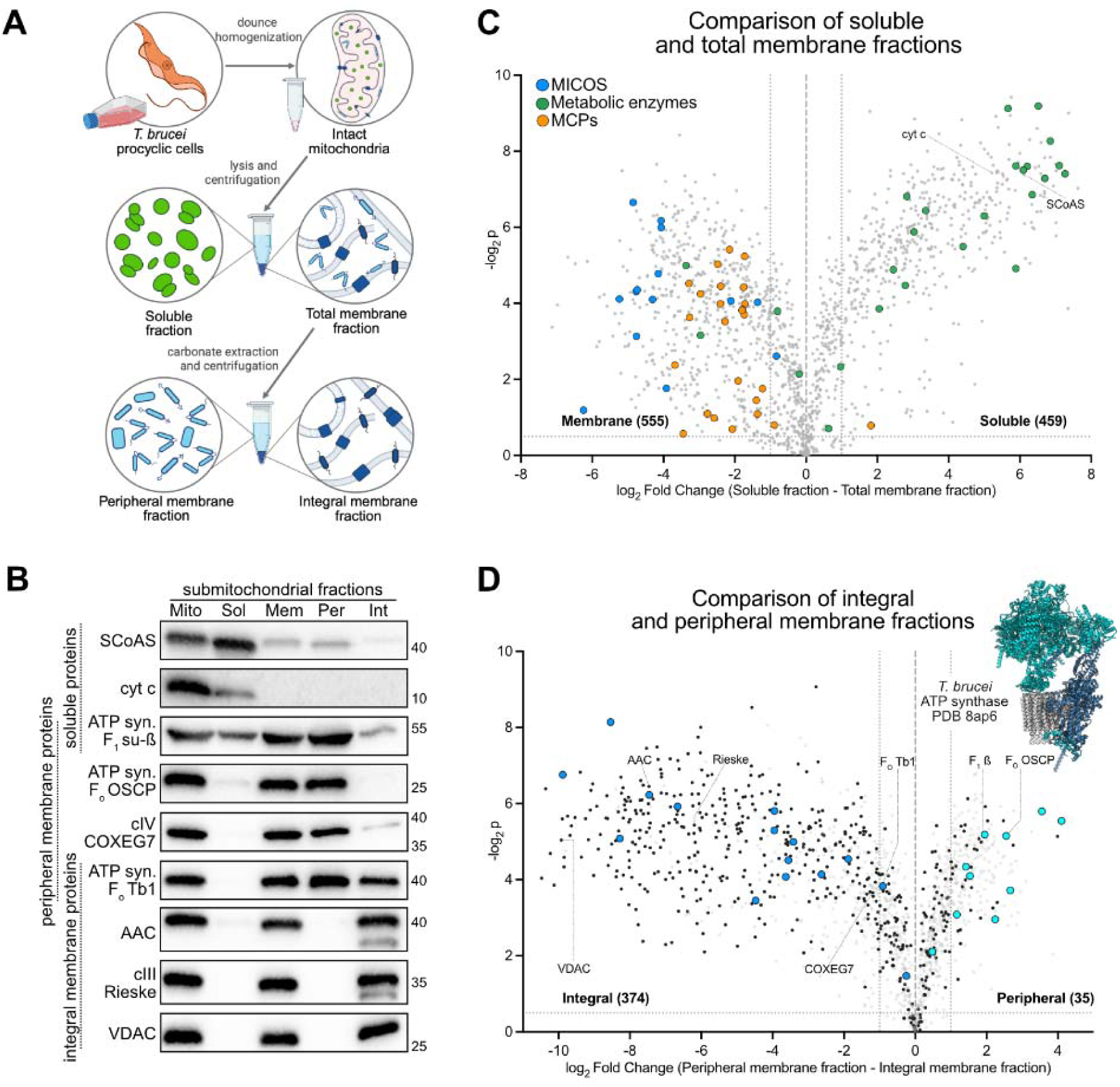
Proteomic characterization of *Trypanosoma brucei* submitochondrial fractions. **(A)** A scheme of fractionation of isolated mitochondria from procyclic trypanosomes. **(B)** Validation of submitochondrial fractionation by immunoblotting of lysates from obtained fractions using indicated antibodies against mitochondrial proteins of known localization (Sol - soluble, Mem - total membrane, Pe - peripheral membrane, Int - integral membrane). **(C)** Validation of the fractionation between soluble and total membrane components (upper panel) and between integral and peripheral membrane components (lower panel) based on the enrichment of indicated mitochondrial proteins and complexes with established localization. MICOS components and MCPs are localized in the membrane fraction and metabolic enzymes are soluble. In the lower panel, ATP synthase subunits with transmembrane helices are shown in blue and peripheral subunits are in cyan. The inset shows the structure of monomeric ATP synthase colored accordingly, with subunits undetectable by MS in white. Dashed lines indicate fold change ±2, and log2p value 0.5.

Overall, we obtained about 90% coverage (1469 proteins for TbOxa1-1 and 1450 proteins for TbOxa1-2 samples) of previously defined mitochondrial proteome of *T. brucei*^32,49,50^. To define proteomes of the submitochondrial fractions, we compared the contents of the soluble and total membrane fractions, and of the peripheral and integral membrane subfractions, in presence of all TbOxa1 homologs. First, to extend the validation of the fractionation procedure beyond the immunoblots described above (Fig. 3B), we analyzed the relative enrichment of the components of the ATP synthase, mitochondrial contact site and cristae organizing system (MICOS) complex, mitochondrial protein import machinery, carrier proteins and selected soluble metabolic enzymes. All these proteins have known submitochondrial localization. The validation confirmed the expected fractionation patterns of most analyzed proteins (Fig. 3C,D). In the case of the ATP synthase, the fractionation of its components between peripheral and integral membrane fractions fully corresponded to its reported structure (^25^; Fig. 3D). Out of mitochondrial-encoded proteins, we detected the cIV subunits COII and COIII, the cIII subunit cytochrome B (CyB), and the product of gene *MURF2*, whose function is unknown. CyB, COII, and COIII were enriched in the integral membrane fraction, consistent with their role as core membrane components of the respective respiratory complexes.

Based on the comparison of soluble and total membrane fractions, the proteins with at least twofold enrichment in soluble or membrane fractions were classified as “*soluble*” and “*membrane*” proteins, respectively (459 and 555 proteins). The remaining 455 proteins were “unclassified”. The *membrane* proteins that showed at least twofold enrichment in the respective fraction based on the peripheral vs integral comparison were further subclassified into “*peripheral*” and “*integral*” proteins (35 and 374 proteins, respectively). 201 out of 374 integral membrane proteins are predicted to contain at least one transmembrane helix, further supporting the quality of separation (Supplementary Table 1).

Our deep-coverage determination of submitochondrial proteomes complements previous proteomic and protein targeting studies on the insect from *T. brucei* mitochondria and its compartments^32,49–52^ and provides valuable datasets for future research of *T. brucei* mitochondrial proteins.

### Submitochondrial proteomic analyses reveal potential substrates of the TbOxa1 machinery

We used the submitochondrial fractionation followed by MS to map the impact of the ablation of TbOxa1-1 and TbOxa1-2 on the proteomes of mitochondrial compartments and to identify the putative substrates of the insertases (Supplementary Table 1). We compared the content of submitochondrial fractions from cells expressing the ectopic copy of the respective paralog, with the state four days after tetracycline removal. We chose time points right before the onset of growth phenotype to minimize impact of secondary effects. TbOxa1-1 and TbOxa1-2 were depleted by 430- and 350-fold, respectively, relative to the control samples, confirming that they are virtually absent when the conditional allele is switched off.

The knockout of TbOxa1-1 resulted in a depletion of 139 proteins from the integral membrane fraction, most of which were classified as integral membrane proteins (see above), while only 25 proteins were enriched (Fig. 4B). The overall trend of membrane protein depletion was also apparent from the comparison of total and peripheral membrane fractions (Supplementary Table 1). Upon the TbOxa1-2 knock-out, depletion of mostly integral membrane proteins from the membrane fractions (Fig. 4E) was accompanied by accumulation of proteins from all classes in the soluble fraction (Fig. 4D). In line with the role of Oxa1 in insertion of hydrophobic mitochondrial encoded OXPHOS subunits into the membrane, a large proportion of the proteins that were most significantly depleted upon loss of TbOxa1-1 and TbOxa1-2 are subunits of electron transport chain complexes or ATP synthase. The absence of TbOxa1-1 resulted predominantly in a depletion of components of complexes cI and cIV, including its mitochondrial encoded subunits COII and COIII, and to a lesser extent cIII (Fig. 4C). In contrast, TbOxa1-2 knockout led to a significant decrease of subunits of cIII, including mitochondrial encoded CyB, and ATP synthase in the membrane fractions (Fig. 4F). In line with this, the components of the matrix-exposed subdomain of ATP synthase accumulated in the soluble fraction in the TbOxa1-2^cKO^ strain (Supplementary Table 1).

**Figure 4.**
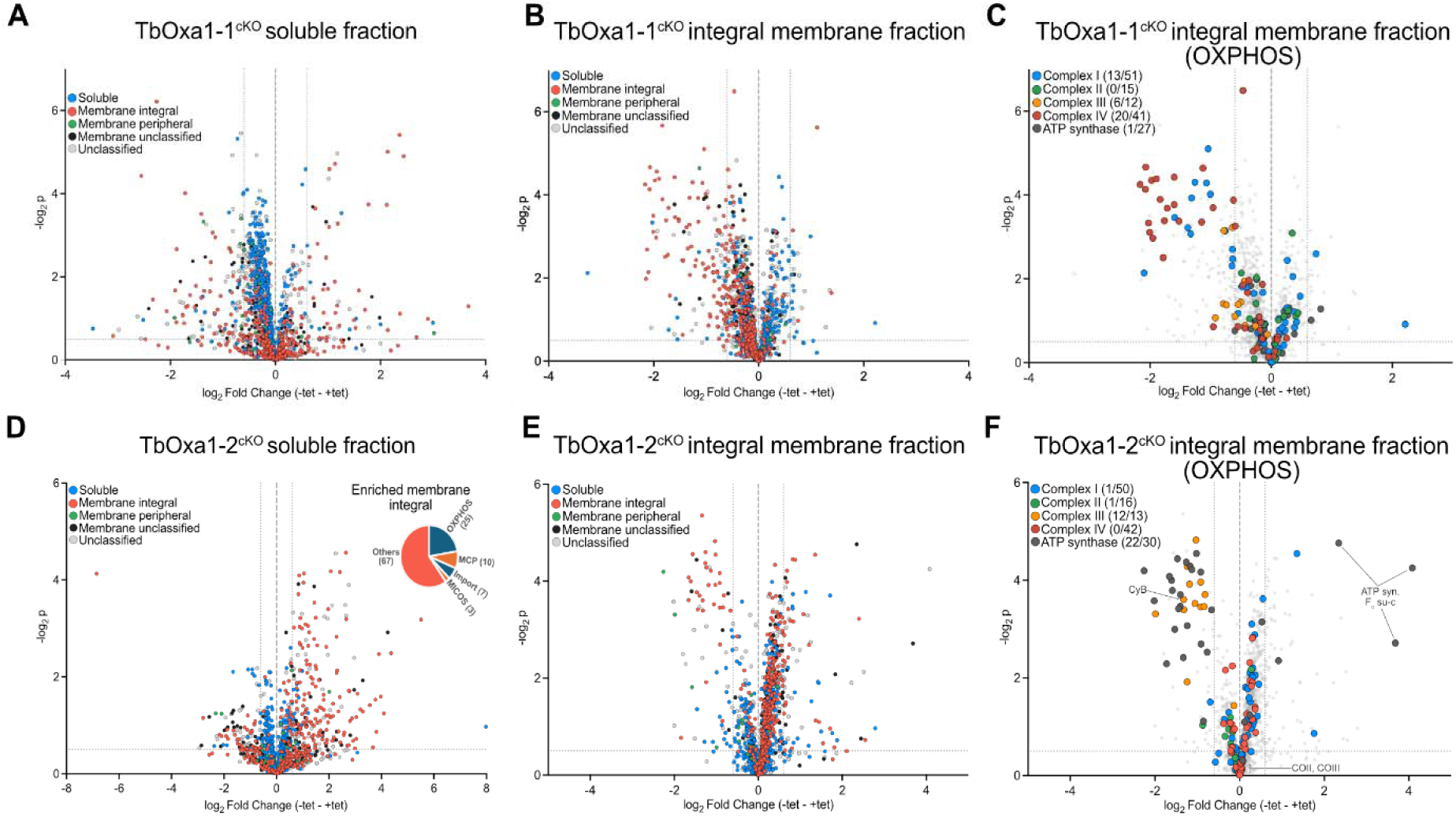
Differential impact of ablation of TbOxa1 paralogs on the content of mitochondrial fractions. (A-F) Dot plots showing the changes in protein content in the soluble (panels (A) and (D)) and integral membrane fractions (panels (B),(C),(E) and (F)) in TbOxa1-1^cKO^ (panels (A) to (C)) and TbOxa1-2^cKO^ (panels (D) to (F)) cells expressing the ectopic copy of the respective gene (+tet) and the same cells 4 (TbOxa1-1^cKO^) or 6 (TbOxa1-2^cKO^ ) days after switching of the ectopic copy by tetracycline removal (-tet). In panels (C) and (F), proteins constituting individual OXPHOS complexes (based on ^46^) are highlighted and the numbers of depleted and total (depleted/total) proteins from each complex are indicated. The dashed lines indicate fold change ±1.5, and log2p value 0.5.

In addition to components of complexes I and IV and several hypothetical proteins of unknown function, the ablation of TbOxa1-1 led to depletion of 12 putative solute carriers from the integral membrane fraction. These included three amino acid transporters, three mitochondrial carrier proteins (MCPs) and other transporters. Furthermore, both paralogs of Mic10, core components of MICOS complex (Kaurov et al., 2018), and two of the four *T. brucei* paralogs of the presequence translocase-associated motor (PAM) J-protein^53^ were also significantly reduced (Supplementary Table 1). A strikingly similar set of proteins was affected by deletion of Oxa1 in *S. cerevisiae*^14,54^, suggesting a conserved function of TbOxa1-1.

In the integral membrane fraction in the TbOxa1-2^cKO^ cells, we observed an enrichment of peptides corresponding to the N-terminal presequences of ATP synthase subunit-c (Fig. 4F). In *T. brucei*, subunit-c is encoded by three nuclear genes; while these isoforms produce identical mature polypeptides found in the ATP synthase complex, they possess distinct presequences. Peptides of the mature subunit-c are undetectable by MS even in the purified enzyme^25^. The ablation of TbOxa1-2 results in 4- to 16-fold accumulation of presequences of all three isoforms. This is in line with the view that TbOxa1-2 is required for the subunit-c insertion into the membrane, documented by the absence of F_1_+c_10_-ring band in native immunoblots in both bloodstream (Fig. 2B) and procyclic (Fig. 2F) trypanosomes. The result shows that the defect in subunit-c insertion is associated with inefficient proteolytic processing by a matrix protease and possibly consequences for import.

Remarkably, numerous integral membrane proteins encoded in the nuclear genome were accumulated in the soluble fraction when TbOxa1-2 was ablated (Fig. 4D), while TbOxa1-1 knock-out resulted in much milder effect (Fig. 4A). Proteins with at least 2-fold accumulation are mostly components of cII to cV OXPHOS complexes, 12 MCPs and five subunits of MICOS (Supplementary Table 1).

We also analysed the overall content of isolated mitochondria to define the TbOxa1-3 depletome. In contrast to pronounced effects of ablation of TbOxa1-1 and TbOxa1-2, only a few mitochondrial proteins were affected in the absence of TbOxa1-3 (3000-fold depletion; Supplementary Fig. 6), in line with the absence of a growth defect (Fig. 2D). The most depleted were mostly membrane-associated hypothetical proteins, mitochondrial carrier protein 23 (TbMCP23), and a cI subunit Tb927.9.15010.

Collectively, the proteomic analysis of mitochondrial compartments upon the inducible knock-out of TbOxa1 paralogs reveals distinct effects: TbOxa1-1 and TbOxa1-2 primarily affect different OXPHOS complexes and integral membrane proteins. Moreover, TbOxa1-2 knock-out results in a pronounced accumulation of membrane proteins, such as MCPs, within the matrix.

### Loss of TbOxa1-1 and TbOxa1-2 is associated with compromised mitochondrial protein synthesis without evidence for a co-translational mechanism

Because insertion of at least some mitochondrial-encoded Oxa1 substrates occurs cotranslationally and Oxa1 directly binds mitoribosomes in human and yeast^36,55^, we asked whether any TbOxa1 paralog(s) interact with mitoribosomes. We chemically crosslinked the content of mitochondria, performed immunoprecipitation (IP) of in situ V5-tagged uL23m and mL78, components of the large mitoribosomal subunit located at the vicinity of the exit from the polypeptide tunnel^56^, and analysed their interacting partners by MS. Despite successful pulldown of mitoribosomes, documented by the enrichment of virtually all protein components, we did not detect any TbOxa1 proteins (Fig. 5A, Supplementary Table 2).

**Figure 5.**
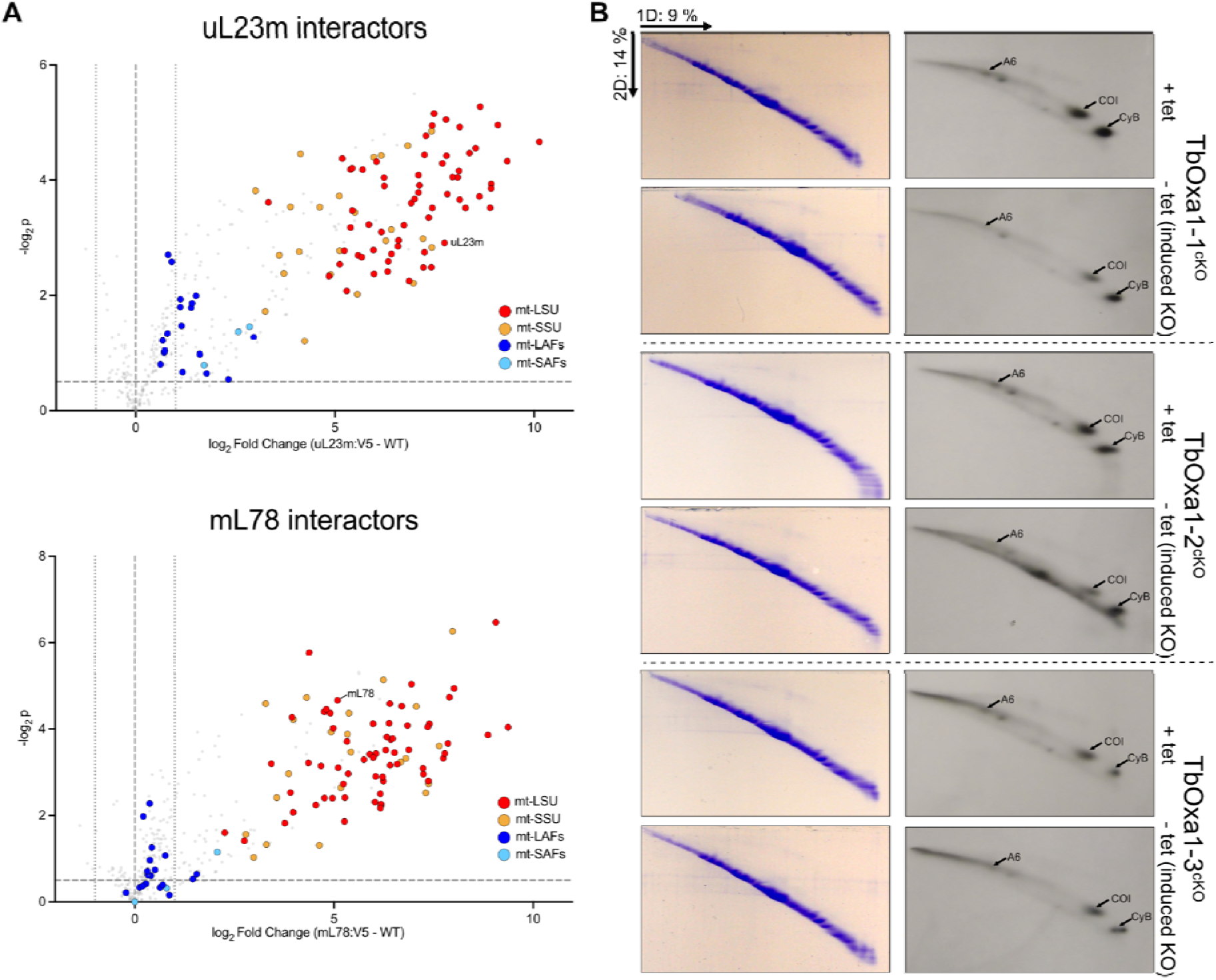
TbOxa1-1 and TbOxa1-2 do not stably interact with mitochondrial ribosomes but are required for efficient mitochondrial translation. **(A)** Dot plots of proteins identified by label-free quantitative MS analysis in samples immunoprecipitated using anti-V5 antibody from cell lines expressing C-terminally V5-tagged mitoribosomal proteins uL23m (top) and mL78 (bottom) as baits compared to control samples from untagged cells. The dashed lines indicate fold change ±1.5, and log2p value 0.5 **(B)** Products of ^35^S metabolic labeling of newly synthesized mitochondrial encoded proteins in TbOxa1^cKO^ cells resolved on 2D SDS-PAGE gels detected by autoradiography (right). Coomassie stained gels (left are shown as loading controls.

In a reciprocal experiment, we immunoprecipitated V5-tagged TbOxa1 paralogs and identified associated proteins by MS (Supplementary Table 2). We detected only three mitoribosomal proteins interacting with TbOxa1-3, while none coimmunoprecipitated with the other two paralogs. In line with these results, *T. brucei* Mba1 - a homolog of a mitoribosome-membrane tethering factor - also failed to pull down mitoribosomal components^31^. Collectively, the IP experiments indicate that the association of mitoribosomes with the membrane insertion machinery in *T. brucei* is weak or transient, if any. However, we note that C-terminal tagging of TbOxa1 paralogs might interfere with their interaction with mitoribosomes.

To directly assay the impact of loss of individual TbOxa1 paralogs on mitochondrial translation, we performed ^35^S metabolic labelling and detected levels of newly synthesized proteins in the conditional knock-out cell lines. In *T. brucei*, this approach requires resolution of labeled products by 2-dimensional denaturing electrophoresis. CyB, COI and subunit a, components of cIII, cIV and ATP synthase, respectively, are the only detectable mitochondrial encoded proteins^57^. While knock-out of TbOxa1-3^cKO^ did not affect translation efficiency, knock-outs of TbOxa1-1 and TbOxa1-2 resulted in a visible decreased signal of all three detectable proteins. While subunit a and COI amounts appeared decreased substantially in both cell lines, the drop of levels of CyB was weaker (Fig. 5B). We did not observe any obvious differences between TbOxa1-1 and TbOxa1-2 knock-outs.

Together, the results show that ablation of both TbOxa1-1 and TbOxa1-2 affect mitochondrial translation to a similar extent, without paralog-specific effects. Our results did not provide evidence for co-translational activity of TbOxa1. Plausibly, the overall decreased translation efficiency can stem from defects in assembly of some or all inner membrane complexes.

### Rhomboid peptidase-like protein acts as a co-factor of TbOxa1-2

Insertases of membrane proteins interact with their substrates and with other components of the membrane biogenesis toolkit. In the IP-MS experiments described above, TbOxa1-1, TbOxa1-2 and TbOxa1-3 co-immunoprecipitated in total 12, 5 and 24 mitochondrial proteins, respectively (1.5-fold enrichment threshold; Fig. 6A; Supplementary Table 2). Interaction partners of TbOxa1 paralogs were mostly OXPHOS components, proteins associated with membrane maintenance, and, rather surprisingly, factors acting in uridine insertion/deletion RNA editing. TbOxa1-2 pulled down Mge1, a nucleotide exchange factor of Hsp70 acting as a component of the PAM complex^58^, reviewed in^59^, in line with a putative role in protein import. TbOxa1-1 was coimmunoprecipitated with TbOxa1-2, suggesting the latter can be a substrate of the former (see Discussion).

**Figure 6.**
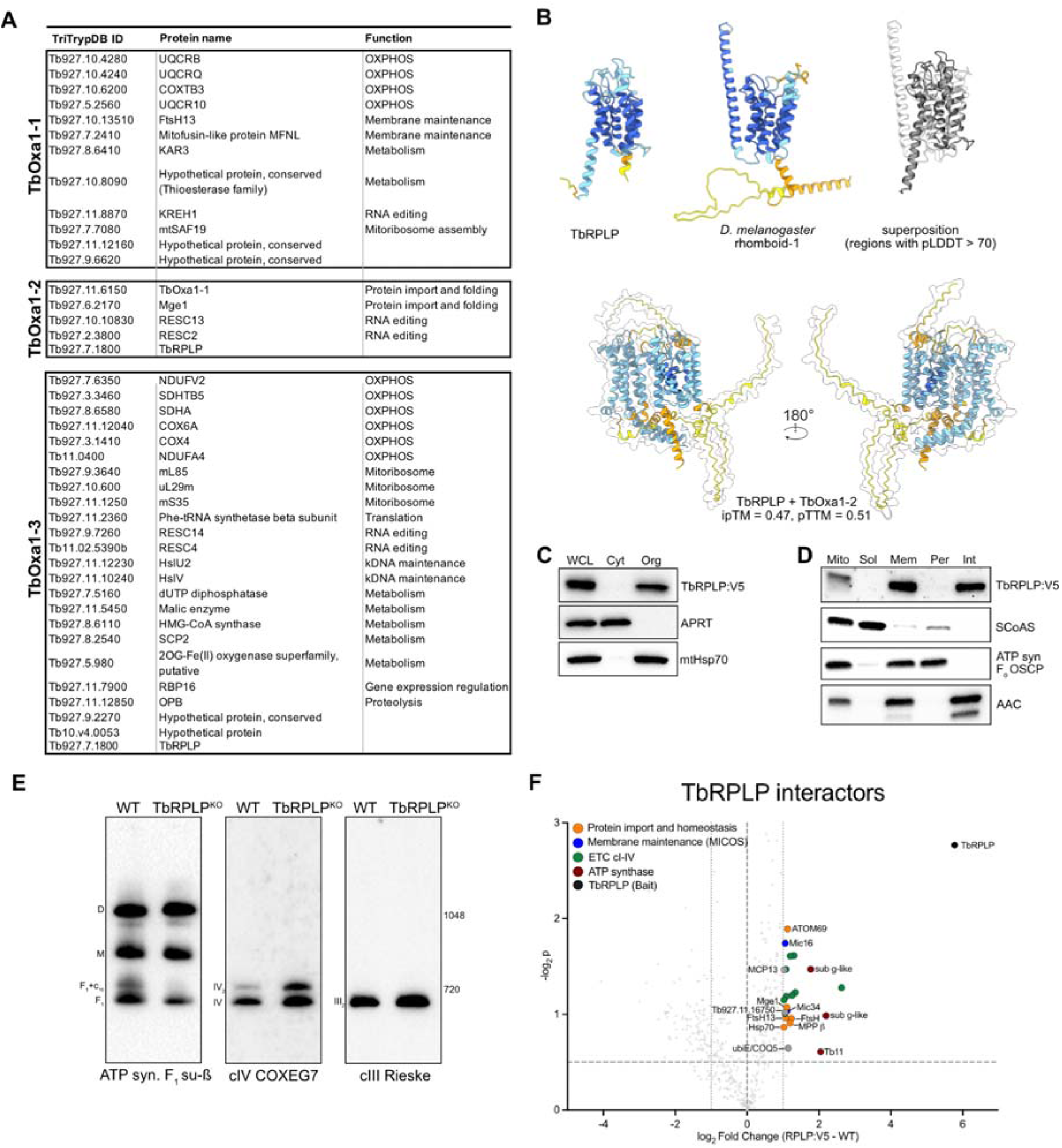
Rhomboid peptidase-like protein acts as a co-factor of TbOxa1-2. **(A)** Mitochondrial proteins that coimmunoprecipitated with V5-tagged TbOxa1 paralogs as baits identified by label-free quantitative MS analysis (fold change ≥ 1.5x, log2p ≥ 0.5). **(B)** Upper panel - AlphaFold models of TbRPLP and rhomboid-1 protease from *Drosophila melanogaster* colored by pLDDT score using the standard AlphaFold palette. In superposition, TbRPLP is shown in grey and rhomboid 1 in white, and regions with pLDDT < 70 are undisplayed. Lower panel - AlphaFold model of a heterodimer of TbRPLP and TbOxa1-2 colored by pLDDT. TbOxa1-2 surface is shown. **(C)** Immunoblots of lysates from whole cells (WCL) and cytosolic (Cyt) and organellar (Org) fractions from V5-tagged TbRPLP-expressing *T. brucei* (TbRPLP:V5) cells. **(D)** Immunoblots of lysates from isolated mitochondria (Mito) and submitochondrial fractions from TbRPLP:V5 cells (Sol - soluble, Mem - total membrane, Per - peripheral membrane, Int - integral membrane). **(E)** Immunoblots of mitochondrial lysates from WT and TbRPLP^KO^ cells resolved by BN-PAGE and probed with indicated antibodies. **(F)** Dot plots of proteins identified by label-free quantitative MS analysis in samples immunoprecipitated using anti-V5 antibody from TbRPLP:V5 cells compared to control samples from untagged cells. The dashed lines indicate fold change ±1.5, and log2p value 0.5 TbRPLP:V5 as bait.

The strongest interaction partner of TbOxa1-2 (14-fold enrichment), and the only protein to pull down with two of the three paralogs, TbOxa1-2 and TbOxa1-3, was a protein annotated in TriTrypDB as hypothetical (Tb927.7.1800). Based on its predicted structure, it is a member of the omnipresent rhomboid-like family (Fig. 6B), which contains membrane-embedded rhomboid peptidases and related proteins without proteolytic activity (Lemberg and Freeman, 2004). In *S. cerevisiae*, the mitochondrial rhomboid peptidase Pcp1 proteolytically processes cytochrome c peroxidase^60^ and membrane remodeler Mgm1 ^61–63^, both localized in the intermembrane space. Deletion of Pcp1 results in the loss of fully assembled ATP synthase assembly and unfolding of cristae^64^. However, the trypanosomal homolog lacks two residues in the catalytic triad essential for the proteolytic activity (N169, S217 and H281 in the prototypic *Drosophila* Rhomboid-1 protease^65^), the N-terminal helix defining mitochondrial rhomboid proteases, and signatures of the metazoan iRhom subfamily of inactive peptidases^66^. Thus, the protein is a highly divergent rhomboid pseudoprotease, which we termed *T. brucei* rhomboid peptidase-like protein (TbRPLP). We modified one allele encoding TbRPLP by fusing it with the V5-epitope tag. The V5-tagged TbRPLP was detected exclusively in the organellar subcellular fraction and in the integral membrane submitochondrial fraction, confirming its localization in the inner mitochondrial membrane (Fig. 6C).

We analyzed the interaction network of TbRPLP by immunoprecipitation with anti-V5 antibody followed by MS (Fig. 6E) and identified 25 mítochondrial proteins with at least 2-fold enrichment, including 17 membrane proteins, out of which 13 are classified as integral (Supplementary Table 1). Although AlphaFold predicts a direct interaction of TbRPLP with TbOxa1-2 (but notably not with TbOxa1-1; Fig. 6B), no TbOxa1 paralog was identified among the interactors. Instead, TbRPLP interacted with 13 OXPHOS-related proteins, including both structural subunits and assembly factors. Two of the strongest interactors are hitherto hypothetical proteins, whose predicted structure revealed they are paralogs of subunit g of ATP synthase (Supplementary Fig. 7A). Subunit g constitutes a dimerization module of ATP synthase complexes^25^. Recently, the two subunit g paralogs were reported to constitute an analogous heterodimeric module connecting an ATP synthase monomer to cIV in a new type of supercomplex^46^. Other interactors include two subunits of MICOS, and five proteins related to protein import, processing and folding, including two putative FtsH-like metalloproteases and PAM complex components Hsp70 and Mge1. Mge1 is a common interactor of TbOxa1-2 and TbRPLP. These results suggest a link between TbRPLP, OXPHOS assembly and import and quality control of proteins.

We knocked-out both alleles encoding TbRPLP by the CRISPR-Cas9 strategy (Supplementary Fig. 7B). Although the knock-out did not result in any growth retardation and steady state levels of OXPHOS subunits and TbOxa1-2 remained largely unchanged (Supplementary Fig. 7C,D), immunoblotting of native OXPHOS complexes revealed the accumulation of high molecular weight cIV-containing assemblies alongside a near-complete loss of the F_1_+c_10_-ring band (Fig. 6E). This phenotype is reminiscent of the defects observed following the ablation of TbOxa1-2 (Fig. 2B,F). Given that the absence of TbOxa1-2 leads to the accumulation of presequences of ATP synthase subunit-c in the membrane (Fig. 4B), we propose that TbOxa1-2 cooperates with TbRPLP and possibly with some of the interacting peptidases to process subunit-c precursors and presumably other membrane proteins.

## DISCUSSION

Insertion of proteins into the inner mitochondrial membrane and assembly of protein complexes in the membrane relies on insertases of bacterial origin from the Oxa1 family. Here, we documented that *Trypanosoma brucei* and related protists from the group Kinetoplastida contain three paralogs of Oxa1, unlike other eukaryotes, which typically possess one or two paralogs. The only other group that contains more than two Oxa1 proteins are diplonemids, free living marine protists related to trypanosomatid parasites. However, Oxa1 duplication occurred independently in kinetoplastids and diplonemids (Fig. 1D & Supplementary Fig. 2). The context and consequences therefore presumably differ in the two groups.

We showed that in *T. brucei*, two of the three paralogs are essential for cultured procyclic form while all three are dispensable for the bloodstream form viability. Knock-out of TbOxa1-1 resulted in pronounced decrease of OXPHOS complexes cI and cIV, while knock-out of TbOxa1-2 affected mostly cIII and ATP synthase. The differential impact on the levels of different complexes is most plausibly explained by the specific role of the two paralogs in insertion of mitochondrially encoded subunits of the respective complexes. Because it is neither possible to metabolically label all 18 mitochondrial encoded proteins in *T. brucei* nor detect them all by mass spectrometry, the substrates of individual TbOxa1 paralogs cannot be determined specifically. However, the absence of apparent paralog-specific effects in the metabolic labeling experiment (Fig. 5B) suggests that depletion of either TbOxa1 paralog does not selectively impair translation of subunits of specific OXPHOS complexes but instead results in a broader reduction in mitochondrial protein synthesis. Nevertheless, individual TbOxa1 paralogs exerted distinct effects on OXPHOS complex biogenesis, evidenced by altered steady-state levels of both mitochondrial and nuclear-encoded subunits of the affected complexes. Analogous functional specialization has been observed in other members of the Oxa1 superfamily. First, in fungi, Oxa2 (or Cox18), a distant Oxa1 paralog, specifically acts in the largely post-translational insertion of Cox2 subunit of cIV, consequently allowing assembly of the entire complex^18–20^. More recently, Oxa2 got duplicated in plants and the resulting paralogs Oxa2A and Oxa2B are required for cytochrome-c maturation and Cox2 insertion, respectively, as documented in *Arabidopsis*^21,67^. Similarly, certain gram-positive bacteria contain two paralogs of YidC, referred to as YidC1 vs YidC2, which display specific functions^68^. Lastly, chloroplast also contain two insertases required for biogenesis of photosystems and other complexes in thylakoids, Alb3 and Alb4^69^, the former of which interacts with the plastid ribosome while the latter does not^70^.

Despite distinct patterns of loss of OXPHOS complexes, we also observed partially overlapping effects upon ablation of TbOxa1-1 and TbOxa1-2. Particularly, cIII and its subunits are depleted not only upon TbOxa1-2 knock-out, but to lesser extent also in TbOxa1-1^KO^ cells (Figs. 2&4). While this can be explained by partial functional redundancy of the two paralogs, there are other possible non-mutually exclusive explanations. Firstly, electron transport chain complexes assemble into supercomplexes (reviewed in^71^). The occurrence of individual complexes in supercomplexes mutually affects their stability and overall abundance. Recently, a new supercomplex consisting of monomeric ATP synthase (cV), cII and two copies of cIV has been reported in *T. brucei*, along with the cIII_2_cIV_2_ supercomplex^46^. The cVcIIcIV_2_ supercomplex caps short rows of ATP synthase dimers, shaping discoidal cristae in trypanosomatid parasites^72,73^. Indeed, cII, which contains no mitochondrial encoded subunit, is slightly decreased in both TbOxa1-1^KO^ and TbOxa1-2^KO^, and high molecular weight complexes containing cIV accumulate in TbOxa1-2^KO^ cells (Fig. 2F). These observations are in accordance with an impact of destabilization of supercomplexes on abundance and stoichiometry of individual complexes. Secondly, because *S. cerevisiae* Oxa1 is required for its own insertion into the membrane^17^, it is reasonable to speculate that in *T. brucei* TbOxa1-1 facilitates insertion of TbOxa1-2. Consequently, the ablation of TbOxa1-1 would result in secondary defects associated with decreased availability of TbOxa1-2. The view that TbOxa1-2 is a substrate of TbOxa1-1 is supported by two observations: (i) TbOxa1-2 is depleted from the membrane fraction upon TbOxa1-1 knock-out and (ii) TbOxa1-1 co-immunoprecipitated with TbOxa1-2 (4-fold enrichment). These results are also in line with the possibility that the two paralogs form active heterodimers. However, although dimerization of Oxa1 was proposed^74^, no direct evidence has been provided and in yeast and mammals Oxa1 associated with mitoribosomes appears to act as monomer, as documented by low to mid resolution cryoEM maps^12,36,75–77^.

Accumulation of membrane proteins in the matrix in absence of TbOxa1-2 suggests a role of this paralog in the conservative sorting of nuclear-encoded proteins. Presumably, these proteins are fully or partially translocated into the matrix, where TbOxa1-2 is required for their re-insertion into the membrane from the inside. This mechanism mirrors the conservative sorting, first documented in *Neurospora crassa*^78^ and later in *S. cerevisiae*^16^. In Oxa1-deficient yeast mutants, this process is compromised and substrate proteins accumulate in the matrix^16^. Alternatively, the proteins are mislocalized to the matrix because they fail to be positioned in the membrane during their transport across the inner mitochondrial membrane due to the absence of TbOxa1-2.

Notably, we observed strong accumulation of presequences of ATP synthase subunit-c in the membrane fraction in TbOxa1-2 knock-out. Subunit-c is a substrate of Oxa1-superfamily insertases in yeast and bacteria^41,42^. The transfer of subunit-c gene to the nuclear genome in trypanosomes implies that, unlike in yeast and bacteria, the insertion is linked to or follows the import. The accumulation of subunit-c presequences in the membrane upon TbOxa1-2 depletion suggests a trapped import intermediate. The mature subunit-c has two transmembrane helices with both termini exposed to the intermembrane space, the topology shared with Cox2, an established Oxa1 substrate budding yeast^79^. The N-terminal presequence is presumably cleaved when exposed to the matrix prior to the insertion of the N-terminal helix to the membrane. The C-terminal helix can be either laterally released into the phospholipid bilayer by the stop transfer mechanism or can be fully imported into the matrix and re-inserted from inside. The absence of TbOxa1-2 results in a failure of the presequence processing. The process might involve newly identified TbRPLP, whose ablation phenocopies the TbOxa1-2 knock-out, and some of the three TbRPLP-interacting proteases. Given the interaction of TbRPLP with subunit g paralogs, which form a module for the integration of ATP synthase into the cVcIIcIV_2_ supercomplex^46^, TbRPLP likely serves as a scaffold or chaperone, which may coordinate the processing of subunit-c precursors with their subsequent insertion by TbOxa1-2.

The expansion of the Oxa1 machinery is consistent with the overall trend of increased proteomic complexity in the *T. brucei* mitochondrion. Gene duplications and protein extension and splitting were shown to contribute to the intricate architecture of trypanosomal mitoribosomes^80,81^, MICOS^82^, or ATP synthase^25,83^. By utilizing three specialized insertases and divergent co-factors like TbRPLP, trypanosomes have developed a robust toolkit to maintain membrane integrity and bioenergetic function in the face of extreme protein divergence. The division of labor between three TbOxa1 proteins may also be an adaptation to specifically manage the insertion and assembly of proteins rendered exceptionally hydrophobic due to extensive uridine insertion editing.

## MATERIALS AND METHODS

### Sequence and structure similarity search, phylogenetic analysis, and structure prediction and analyses

Basic sequence similarity search was performed using blastp^84^. Structure-based similarity was detected by FoldSeek^85^. Structure prediction was conducted using AlphaFold 3^86^ and transmembrane helices were predicted by DeepTMHMM 1.0^87^. Structures were depicted using ChimeraX^88^.

For the phylogenetic analyses, initial sequences of putative Oxa1 proteins were retrieved from predicted proteomes of all organisms from the group Euglenozoa in EukProt v3 database^89^ by blastp search using *T. brucei* TbOxa1-1, TbOxa1-2 and TbOxa1-3 as queries. Sequences of Oxa1 proteins classified as “reviewed” in the Uniprot database were added, and misannotated proteins were identified by CLANS clustering analysis^90^ and removed. The resulting sequences were aligned using MAFFT v7.525^91^ (L-INS-i algorithm) and from the alignment, a profile hmm was built in HMMER3^92^. The profile was used to search predicted proteomes of selected 29 organisms from the group Euglenozoa in the EukProt v3 database. Bacterial contaminants and false positive hits were removed based on CLANS clustering analysis, SWISS-MODEL^93^ predictions and preliminary phylogenetic analysis with parameters described below. For the final phylogenetic tree, the dataset was supplemented by sequences of reference eukaryotic Oxa1 and Alb3 proteins, and bacterial YidC from both UniProt and NCBI databases. Sequences were aligned as described above and trimmed using BMGE v1.12^94^. The maximum likelihood tree was inferred with IQ-TREE^95^ multicore v2.3.5 using LG+C20+G4 model. Statistical support was assessed by 1000 ultrafast bootstraps^96^ with bnni correction and SH-alrt test^97^. The phylogenetic tree was rooted between bacterial YidC and eukaryotic Oxa1 and visualized using FigTree v1.4.4.

### Cell culture and generation of cell lines

All procyclic form and bloodstream form *T. brucei* strains in this study were derived from the respective form of the Lister 427 strain. Procyclic cells were cultured at 27°C in SDM-79 medium supplemented with 10% (v/v) fetal bovine serum (FBS). Bloodstream form cells were grown at 37°C in HMI-11 medium supplemented with 10% FBS. Conditional knockout procyclic cells were grown in the presence of hygromycin (25 μg/ml), neomycin (15 μg/ml), phleomycin (2.5 ug/ml) and puromycin (1 μg/ml), depending on the respective selection marker. V5 epitope-tagged cells were cultivated in the presence of hygromycin (5 μg/ml) and puromycin (1 μg/ml).

To express C-terminally V5 epitope-tagged TbOxa1 proteins, DNA fragments containing the epitope and selection marker cassette were generated by PCR using pPOTv5 plasmid as template and primers (Table S3) with 80-nucleotide long extensions for homologous recombination at the 3’ end of the gene of interest upstream the stop codon (Dean et al., 2015). The PCR products were transfected into *T. brucei* procyclic cells, clones resistant to selection antibiotics were selected and screened by sodium dodecyl sulfate (SDS) polyacrylamide gel electrophoresis (PAGE) and immunoblotting of whole cell lysates with anti-V5 antibody.

All knock-out cell lines were generated in cells constitutively expressing T7 RNA polymerase, Tet repressor and Cas9 nuclease from a single construct (procyclic or bloodstream form SmOx Cas9 cells)^98^. For the conditional expression of TbOxa1 proteins in procyclic cells, an ectopic tetracycline regulatable copy of the respective gene was introduced to the parental strain by transfection of a *Not*I-linearized pLEW79-based construct carrying the respective coding sequence (CDS) inserted between the *Bam*HI and *Hind*III sites and the phleomycin resistance marker. Positive clones were selected using phleomycin (2.5 μg/ml). To knockout both alleles of individual genes of interests in procyclic cells, two linear DNA molecules for expression of single guide RNAs (sgRNAs) and two repair templates containing selection marker genes with UTRs flanked by homology arms targeting their recombination to the locus of interest were generated by PCR with oligonucleotides designed using the LeishGEdit tool^99^ (Supplementary Fig. 5A). The repair templates were amplified from donor plasmids pPOTv5 (hygromycin) and pHD2164 (neomycin). The cells were electroporated with a mixture of both sgRNAs and both repair templates (4 μg each) using Amaxa Nucleofactor II (program X-014). Clonal populations of knockout cells were selected in SDM-79 medium supplemented with hygromycin (25 μg/ml) and neomycin (15 μg/ml).

To generate gene knockouts in bloodstream form SmOx Cas9 cells, two linear DNA molecules for expression of sgRNAs and a single repair template for expression of selection marker genes with UTRs flanked by homology arms (4 μg each) were introduced into the cells by electroporation using Amaxa Nucleofactor II (program X-001; Supplementary Fig. 4A). The repair templates were amplified from donor plasmids pPOTv5 (hygromycin), pAZ0055 (phleomycin) and pHD2164 (neomycin). Clonal populations of knockout cells were selected in HMI-11 medium supplemented with hygromycin (5 μg/ml), phleomycin (2.5 μg/ml), or neomycin (2.5 μg/ml).

In selected clones, the deletion of CDS of genes of interest and replacement by selection marker genes was verified by PCR with specific primers (Supplementary Figs. S4B & S5B). The sequences of all oligonucleotides are provided in Table S3.

### Mitochondrial isolation

Mitochondria were isolated from 2.5×10^8^ *T. brucei* cells in the exponential growth. The cells were harvested by centrifugation (1300*g*, 10 minutes, room temperature), washed in PBS-G buffer (20 mM sodium phosphate buffer pH 7.9 with 150 mM NaCl and 20 mM glucose), resuspended in hypotonic buffer (1mM Tris-HCl pH 8.0, 1mM EDTA), and ruptured in a Dounce homogenizer with 10 strokes. Sucrose was immediately added to a final concentration of 0.25 M to restore isotonic conditions. Crude mitochondria were pelleted (16000*g*, 10 minutes, 4°C), resuspended in STM buffer (250 mM sucrose, 20 mM Tris-HCl pH 8.0, 5 mM MgCl_2_, 0.3 mM CaCl_2_ and treated with 5 ug/ml DNase I. After incubation on ice for 1 hour, one volume of STE buffer (20 mM Tris-HCl pH 8.0, 250 mM sucrose, 2 mM EDTA) was added, and mitochondria were pelleted (16000*g*, 10 minutes, 4°C). The pellets were snap-frozen in liquid nitrogen and stored at -80°C.

### Subcellular and submitochondrial fractionation

To prepare subcellular fractions, 1×10^8^ cells were harvested (1300*g*, 10 minutes, room temperature), washed in cold PBS-G, and spun down (1300*g*, 10 minutes, 4°C). The cell pellets were resuspended and lysed in SoTE buffer (0.6 M sorbitol, 20 mM Tris-HCl pH 7.5, 2 mM EDTA) supplemented with 0.015% (w/v) digitonin. After incubation on ice for 5 minutes, the organellar pellet was separated from the cytosolic fraction (supernatant) by centrifugation (4600*g*, 3 minutes, 4°C). The organellar pellet was resuspended in the same volume of PBS buffer (20 mM sodium phosphate buffer pH 7.9 with 150 mM NaCl). Both fractions were treated with SDS-PAGE dye (150 mM Tris pH 6.8, 300 mM 1,4-dithiothreitol, 6% (w/v) SDS, 30% (w/v) glycerol, 0.02% (w/v) bromophenol blue), heated at 97°C for 7 minutes, and used for electrophoresis and immunoblotting.

For mitochondrial subfractionation, hypotonically isolated mitochondria from 1.5×10^9^ procyclic *T. brucei* cells were resuspended in 10 mM MgCl_2_ and was subjected to 10 successive cycles of flash freezing in liquid nitrogen and thawing at room temperature. After centrifugation (6000*g*, 5 minutes, 4°C), the supernatant (soluble matrix fraction) was separated from the pellet (total membrane fraction). A fraction of the pellet was resuspended in 0.1 M Na_2_CO_3_ pH 11.5, transferred to polypropylene tubes, and subjected to ultracentrifugation (100,000*g*) in SW50Ti rotor for 23 minutes at 4°C. The supernatant (peripheral membrane protein fraction) was collected, and the pellet (integral membrane protein fraction) was resuspended in 10 mM MgCl_2_. The individual fractions were analyzed by electrophoresis and immunoblotting, and by label-free quantitative mass spectrometry.

### Proteinase K protection assay

Proteinase K protection assay was performed according to Kaurov et al., 2018^37^. Briefly, mitochondria-enriched organellar fraction from 1×10^8^ *T. brucei* procyclic cells (see subcellular fractionation protocol described above) was resuspended in SoTE buffer (0.6 M sorbitol, 20 mM Tris-HCl pH 7.5, 2 mM EDTA) with 0.03% digitonin. After incubation on ice for 10 minutes, the pellet (“mitoplast”) was collected by centrifugation (4600*g*, 3 minutes, 4°C), resuspended in SoTE buffer, and split into three equal volumes. Two aliquots were treated with 0.5 mg/mL of proteinase K, and one of them was further treated with 1% Triton-X (v/v). After incubation on ice for 30 minutes, the reactions were stopped by the addition of 5 mM PMSF.

### Quantitative polymerase chain reaction

Isolation of total RNA from 5×10^7^ *T. brucei* cells was performed using the MasterPure Complete DNA and RNA Purification Kit (Lucigen MC85200) following the manufacturer protocol. The concentration and quality of isolated RNA were assessed by microvolume spectrophotometer and electrophoretically. Synthesis of cDNA from isolated RNA samples was carried out using TaqMan Reverse Transcription kit (Applied Biosystems N8080234) following the manufacturer protocol. The synthesized cDNA samples were subsequently subjected to quantitative PCR using the LightCycler 480 SYBR Green I Master (Roche 04707516001) on the LightCycler 480 Instrument. The oligonucleotides employed for detection of cDNA targets are listed in Supplementary Table 3.

### Immunoprecipitation

Dynabeads Protein G (Invitrogen 10003D) were conjugated to monoclonal anti-V5 antibody by crosslinking with 11 mM dimethyl suberimidate in PBS buffer pH 8.0 for 30 minutes at room temperature. The crosslinking reaction was quenched by the addition of 50 mM Tris-HCl pH 7.5, washed with PBS buffer with 0.1% (v/v) Tween20, and finally with IP buffer (25 mM Tris-HCl pH 7.5, 100 mM KCl, 15mM MgCl_2_, 0.1% (v/v) n-dodecyl -D-maltoside, EDTA-free protease inhibitor cocktail). Hypotonically isolated mitochondria from 5×10^8^ *T. brucei* procyclic cells were resuspended in PBS pH 7.7 and chemically crosslinked using 1 mM Lomant’s reagent (Thermo Scientific Pierce 22586). After incubation for 2 hours on ice, the reaction was stopped by the addition of 20 mM Tris-HCl pH 7.5. The pellet was collected by centrifugation (16000*g*, 15 minutes, 4°C), resuspended in the IP buffer with 1% (v/v) DDM, and incubated on ice for 1 hour. The samples were spun down (16000*g*, 1 hour, 4°C). The resulting lysates were collected and incubated with V5 antibody-bound Dynabeads for 2 hours at 4°C. The lysates (unbound fraction) were transferred to a fresh tube, and the beads were washed twice with the IP buffer. The beads were snap-frozen, stored at -80°C, and submitted for label-free mass spectrometry.

### Mass spectrometry analyses

Submitochondrial fractions and samples from immunoprecipitations, each in three replicates, were analyzed by label-free quantification mass spectrometry analysis (LFQ-MS) following the SP4 no glass bead protocol (Solvent Precipitation SP3 (SP4)^100^). IP samples were processed similarly with the addition of glass beads. Briefly, proteins were solubilized by 1% SDS in 100 mM triethylammonium bicarbonate (TEAB) buffer, reduced with 10 mM Tris(2-carboxyethyl)phosphine (TCEP), and alkylated with 40 mM chloroacetamide (performed together at 95°C for 10 min). Proteins were digested overnight at 37°C with MS-grade trypsin at a 1:40 enzyme-to-protein ratio. The resulting peptides were desalted using C18 StageTips. Approximately 500 ng of peptide digest per sample was separated on a C18 column on nanoUHPLC Dionex Ultimate 3000 and analyzed in data-independent acquisition mode (DIA) on an

Orbitrap Exploris 480 mass spectrometer equipped with a FAIMS unit. IP samples were acquired in a DDA mode and analyzed in MaxQuant. DIA MS Thermo raw files were processed in Spectronaut (Biognosys) using directDIA mode and *Trypanosoma brucei* proteomeTriTrypDB-68_TbruceiTREU927_AnnotatedProteins.fasta and default setting with Precursor and Protein Q-value and PEP cutoff set at 0.01. Protein group quantities (PG.Quantity, MS2 level) from Spectronaut’s protein report were analyzed in Perseus software.

### Production of polyclonal antibody against TbOxa1-2

The coding sequence of TbOxa1-2 was PCR amplified from genomic DNA. The reverse primer contained sequence encoding hexahistidine (6xHis) tag (Supplementary Table 3). The PCR product was cloned into the expression vector pSKB3 and the resulting construct was transformed into the BL21-CodonPlus(DE3)-RIPL *Escherichia coli* strain. The protein expression was induced with 1 mM IPTG for 3 hours. The heterologously expressed TbOxa1-2-6xHis protein was solubilized in the presence of 2 % sarkosyl, purified by nickel affinity chromatography and dialyzed to a buffer containing 0.7 % sarkosyl. The purified protein was used as an antigen for polyclonal antibody production (standard immunization protocol, Davids Biotechnology, Regensburg, Germany).

### Protein electrophoresis, immunoblotting, and in-gel activity assays

Denaturing and blue native PAGE and immunoblotting were performed as described earlier^25^. Briefly, whole cell lysates for denaturing SDS-PAGE were resuspended in PBS buffer and 3x Laemmli buffer (150 mM Tris pH 6.8, 300 mM 1,4-dithiothreitol, 6% (w/v) SDS, 30% (w/v) glycerol, 0.02% (w/v) bromophenol blue) to a final concentration of 1×10^5^ cells/μL. The lysates were incubated at 97°C for 7 minutes and used for electrophoresis. Sample corresponding to material from 1×10^6^ cells were separated on 4-20% gradient Tris-glycine polyacrylamide gels (BioRad 4568094), transferred onto a PVDF membrane (Pierce 88518), and immunoprobed with respective antibodies (Supplementary Table 3). The membranes were incubated with the Clarity Western ECL substrate (BioRad 1705060EM) and chemiluminescence was detected on a ChemiDoc instrument (BioRad).

For blue native electrophoresis, crude mitochondrial vesicles from 2.5×10^8^ cells were resuspended in 40 μl of solubilization buffer A (2 mM E-aminocaproic acid (ACA), 1 mM EDTA, 50 mM NaCl, 50 mM Bis-Tris/HCl pH 7.0) and solubilized with 2% DDM (w/v) for 1 hour on ice. Lysates were collected by centrifugation (16000*g*, 1 hour, 4°C) and their protein concentration was determined colorimetrically using bicinchoninic acid (BCA) assay. For each sample, 4 μg of total protein from mitochondrial lysates was mixed with 1.5 μl of loading dye (500 mM ACA, 5% (w/v) Coomassie Brilliant Blue G-250, 5% (w/v) glycerol) to a final volume of 20 μl/well. The samples were resolved on a 3-12% Bis-Tris native PAGE gel (Invitrogen). After electrophoretic separation (3 hours, 140 V, 4°C), the proteins were transferred onto a PVDF membrane by electroblotting (100 minutes, 100 V, 4°C), followed by immunodetection with appropriate antibodies (Supplementary Table 3). In-gel staining of activity of complex IV was performed by incubating BN-PAGE gels in assay buffer (50 mM phosphate buffer pH 7.4, 1 mg/mL 3,3′diaminobenzidine, 24 U/mL catalase, 1 mg/mL cytochrome *c*, 75 mg/mL sucrose) with slow agitation overnight.

### Membrane potential measurement

Spectrofluorometric assessment of mitochondrial membrane polarization in digitonin-permeabilized *T. brucei* bloodstream form cells using safranin O dye (Sigma S2255) was performed as described previously ^101^ with some modifications. Briefly, 2×10^7^ cells were pelleted (1300*g*, 10 minutes, room temperature) and washed with 1 ml of ANT buffer (8 mM KCl, 110 nM K-gluconate, 10 mM NaCl, 10 mM free-acid Hepes, 10 mM K_2_HPO_4_, 0.015 mM EGTA potassium salt, 10 mM mannitol, 0.5 ml/ml fatty acid-free BSA, 1.5 mM MgCl_2_, pH 7.25). The cells were permeabilized by 4 μM digitonin (Sigma D141) in 2 ml of ANT buffer with 5 μM safranin O. Fluorescence was recorded for 750 s in Hitachi F-7000 spectrofluorimeter (Hitachi High Technologies) with the following parameters: 2 Hz acquisition rate and 495 nm and 585 nm excitation and emission wavelengths, respectively. 1 mM ATP and 10 μg/ml oligomycin were added after 250 s and 600 s, respectively. The uncoupler SF 6847 (250 nM; Enzo Life Sciences BML-EI215-0050) was added to trigger maximal depolarization.

### Metabolic labeling of mitochondrial translation products and two-dimensional electrophoresis

For labeling of products of mitochondrial translation, 10^7^ cells were harvested from the culture in logarithmic phase of growth (5 minutes, 1000 x *g*, RT), washed twice in SoTE medium (0.6 M sorbitol, 20 mM Tris-HCl, pH 7.5, 2 mM EDTA), pelleted (5 minutes, 1100 x *g*, RT) and finally homogenized in 90 μl of SoTE. Cycloheximide and dithiothreitol were added to the cell suspension to final concentration of 200 μg/ml and 2 mM, respectively, to inhibit cytosolic translation and prevent cells from forming aggregates. After 10 minutes of incubation at 27 °C with vigorous shaking, the products of mitochondrial translation were labeled *in vivo* for 1 hour using 10 μl of Met-^35^S-Label mix (Hartmann Analytic) per sample. Labeled products were resolved in two-dimensional Tris-glycine-SDS PAGE (9% in the first dimension, 14% in the second dimension) and visualized by autoradiography as described previously^102^. The total proteins were visualized with Coomassie brilliant blue R250 staining.

## DATA AVAILABILITY

All data needed to evaluate conclusions of this study are available in the manuscript or as Supplementary Files. The mass spectrometry proteomic data have been deposited to the PRIDE repository under accession numbers XXX and XXX. Source data are provided with this manuscript.

## ACKNOWLEDGEMENTS

We thank Marek Vrbacký for processing and analyses of mass spectrometry proteomics samples and deposition of datasets, and Carolina Hierro-Yap for help with several experiments.

## AUTHOR CONTRIBUTIONS

Conceptualization: J.E.W., O.G. Methodology: J.E.W., I.Š.S., A.Z., O.G. Formal Analysis: J.E.W., J.Ř. Investigation: J.E.W, I.Š.S., J.Ř., P.C., A.L., V.D. Resources: A.Z. Writing – Original Draft: J.E.W., O.G. Writing – Review & Editing: J.E.W., A.Z., O.G. Visualization: J.E.W., J.Ř., O.G. Funding Acquisition: J.E.W., I.Š.S., P.C., A.Z., O.G.

## REFERENCES

1 Bonnefoy, N. et al. Cloning of a human gene involved in cytochrome oxidase assembly by functional complementation of an oxa1- mutation in *Saccharomyces cerevisiae*. Proc Natl Acad Sci U S A 91, 11978–11982, doi:10.1073/pnas.91.25.11978 (1994).

2 Samuelson, J. C. et al. YidC mediates membrane protein insertion in bacteria. Nature 406, 637–641, doi:10.1038/35020586 (2000).

3 Scotti, P. A. et al. YidC, the *Escherichia coli* homologue of mitochondrial Oxa1p, is a component of the Sec translocase. EMBO J 19, 542–549, doi:10.1093/emboj/19.4.542 (2000).

4 McDowell, M. A., Heimes, M. & Sinning, I. Structural and molecular mechanisms for membrane protein biogenesis by the Oxa1 superfamily. Nat Struct Mol Biol 28, 234–239, doi:10.1038/s41594-021-00567-9 (2021).

5 Pyrih, J. et al. Vestiges of the Bacterial Signal Recognition Particle-Based Protein Targeting in Mitochondria. Mol Biol Evol 38, 3170–3187, doi:10.1093/molbev/msab090 (2021).

6 He, S. & Fox, T. D. Membrane translocation of mitochondrially coded Cox2p: distinct requirements for export of N and C termini and dependence on the conserved protein Oxa1p. Mol Biol Cell 8, 1449–1460, doi:10.1091/mbc.8.8.1449 (1997).

7 Hennon, S. W., Soman, R., Zhu, L. & Dalbey, R. E. YidC/Alb3/Oxa1 Family of Insertases. J Biol Chem 290, 14866–14874, doi:10.1074/jbc.R115.638171 (2015).

8 Homberg, B., Rehling, P. & Cruz-Zaragoza, L. D. The multifaceted mitochondrial OXA insertase. Trends Cell Biol, doi:10.1016/j.tcb.2023.02.001 (2023).

9 Hell, K., Neupert, W. & Stuart, R. A. Oxa1p acts as a general membrane insertion machinery for proteins encoded by mitochondrial DNA. EMBO J 20, 1281–1288, doi:10.1093/emboj/20.6.1281 (2001).

10 Bonnefoy, N., Fiumera, H. L., Dujardin, G. & Fox, T. D. Roles of Oxa1-related inner-membrane translocases in assembly of respiratory chain complexes. Biochim Biophys Acta 1793, 60–70, doi:10.1016/j.bbamcr.2008.05.004 (2009).

11 Ott, M. & Herrmann, J. M. Co-translational membrane insertion of mitochondrially encoded proteins. Biochim Biophys Acta 1803, 767–775, doi:10.1016/j.bbamcr.2009.11.010 (2010).

12 Schondorf, T. et al. Membrane insertion of mitochondrial-encoded proteins regulates ribosome decoding speed. Nat Struct Mol Biol 33, 853–867, doi:10.1038/s41594-026-01803-w (2026).

13 Neupert, W. & Herrmann, J. M. Translocation of proteins into mitochondria. Annu Rev Biochem 76, 723–749, doi:10.1146/annurev.biochem.76.052705.163409 (2007).

14 Stiller, S. B. et al. Mitochondrial OXA Translocase Plays a Major Role in Biogenesis of Inner-Membrane Proteins. Cell Metab 23, 901–908, doi:10.1016/j.cmet.2016.04.005 (2016).

15 Bohnert, M. et al. Cooperation of stop-transfer and conservative sorting mechanisms in mitochondrial protein transport. Curr Biol 20, 1227–1232, doi:10.1016/j.cub.2010.05.058 (2010).

16 Hell, K., Herrmann, J. M., Pratje, E., Neupert, W. & Stuart, R. A. Oxa1p, an essential component of the N-tail protein export machinery in mitochondria. Proc Natl Acad Sci U S A 95, 2250–2255, doi:10.1073/pnas.95.5.2250 (1998).

17 Herrmann, J. M., Neupert, W. & Stuart, R. A. Insertion into the mitochondrial inner membrane of a polytopic protein, the nuclear-encoded Oxa1p. EMBO J 16, 2217–2226, doi:10.1093/emboj/16.9.2217 (1997).

18 Funes, S., Nargang, F. E., Neupert, W. & Herrmann, J. M. The Oxa2 protein of *Neurospora crassa* plays a critical role in the biogenesis of cytochrome oxidase and defines a ubiquitous subbranch of the Oxa1/YidC/Alb3 protein family. Mol Biol Cell 15, 1853–1861, doi:10.1091/mbc.e03-11-0789 (2004).

19 Souza, R. L., Green-Willms, N. S., Fox, T. D., Tzagoloff, A. & Nobrega, F. G. Cloning and characterization of COX18, a *Saccharomyces cerevisiae* PET gene required for the assembly of cytochrome oxidase. J Biol Chem 275, 14898–14902, doi:10.1074/jbc.275.20.14898 (2000).

20 Saracco, S. A. & Fox, T. D. Cox18p is required for export of the mitochondrially encoded *Saccharomyces cerevisiae* Cox2p C-tail and interacts with Pnt1p and Mss2p in the inner membrane. Mol Biol Cell 13, 1122–1131, doi:10.1091/mbc.01-12-0580 (2002).

21 Kolli, R., Soll, J. & Carrie, C. OXA2b is Crucial for Proper Membrane Insertion of COX2 during Biogenesis of Complex IV in Plant Mitochondria. Plant Physiol 179, 601–615, doi:10.1104/pp.18.01286 (2019).

22 Maslov, D. A. et al. Recent advances in trypanosomatid research: genome organization, expression, metabolism, taxonomy and evolution. Parasitology 146, 1–27, doi:10.1017/S0031182018000951 (2019).

23 Acestor, N., et al. *Trypanosoma brucei* mitochondrial respiratome: composition and organization in procyclic form. Mol Cell Proteomics 10, M110 006908, doi:10.1074/mcp.M110.006908 (2011).

24 Schneider, A. Evolution of mitochondrial protein import - lessons from trypanosomes. Biol Chem 401, 663–676, doi:10.1515/hsz-2019-0444 (2020).

25 Gahura, O. et al. An ancestral interaction module promotes oligomerization in divergent mitochondrial ATP synthases. Nat Commun 13, 5989, doi:10.1038/s41467-022-33588-z (2022).

26 Gahura, O., Hierro-Yap, C. & Zikova, A. Redesigned and reversed: architectural and functional oddities of the trypanosomal ATP synthase. Parasitology 148, 1151–1160, doi:10.1017/S0031182021000202 (2021).

27 Aphasizheva, I. et al. Lexis and Grammar of Mitochondrial RNA Processing in Trypanosomes. Trends Parasitol 36, 337–355, doi:10.1016/j.pt.2020.01.006 (2020).

28 Moller-Hergt, B. V., Carlstrom, A., Stephan, K., Imhof, A. & Ott, M. The ribosome receptors Mrx15 and Mba1 jointly organize cotranslational insertion and protein biogenesis in mitochondria. Mol Biol Cell 29, 2386–2396, doi:10.1091/mbc.E18-04-0227 (2018).

29 Preuss, M. et al. Mba1, a Novel Component of the Mitochondrial Protein Export Machinery of the Yeast*Saccharomyces cerevisiae*. The Journal of Cell Biology 153, 1085–1096, doi:10.1083/jcb.153.5.1085 (2001).

30 Ott, M. et al. Mba1, a membrane-associated ribosome receptor in mitochondria. EMBO J 25, 1603–1610, doi:10.1038/sj.emboj.7601070 (2006).

31 Wenger, C. et al. The Mba1 homologue of *Trypanosoma brucei* is involved in the biogenesis of oxidative phosphorylation complexes. Mol Microbiol, doi:10.1111/mmi.15048 (2023).

32 Pyrih, J. et al. Comprehensive sub-mitochondrial protein map of the parasitic protist *Trypanosoma brucei* defines critical features of organellar biology. Cell Rep 42, 113083, doi:10.1016/j.celrep.2023.113083 (2023).

33 Kumazaki, K. et al. Structural basis of Sec-independent membrane protein insertion by YidC. Nature 509, 516–520, doi:10.1038/nature13167 (2014).

34 Kumazaki, K. et al. Crystal structure of *Escherichia coli* YidC, a membrane protein chaperone and insertase. Sci Rep 4, 7299, doi:10.1038/srep07299 (2014).

35 Szyrach, G., Ott, M., Bonnefoy, N., Neupert, W. & Herrmann, J. M. Ribosome binding to the Oxa1 complex facilitates co-translational protein insertion in mitochondria. EMBO J 22, 6448–6457, doi:10.1093/emboj/cdg623 (2003).

36 Itoh, Y. et al. Mechanism of membrane-tethered mitochondrial protein synthesis. Science 371, 846–849, doi:10.1126/science.abe0763 (2021).

37 Kaurov, I. et al. The Diverged Trypanosome MICOS Complex as a Hub for Mitochondrial Cristae Shaping and Protein Import. Curr Biol 28, 3393–3407 e3395, doi:10.1016/j.cub.2018.09.008 (2018).

38 Nolan, D. P. & Voorheis, H. P. The mitochondrion in bloodstream forms of *Trypanosoma brucei* is energized by the electrogenic pumping of protons catalysed by the F1F0-ATPase. Eur J Biochem 209, 207–216, doi:10.1111/j.1432-1033.1992.tb17278.x (1992).

39 Schnaufer, A., Clark-Walker, G. D., Steinberg, A. G. & Stuart, K. The F1-ATP synthase complex in bloodstream stage trypanosomes has an unusual and essential function. EMBO J 24, 4029–4040, doi:10.1038/sj.emboj.7600862 (2005).

40 Brown, S. V., Hosking, P., Li, J. & Williams, N. ATP synthase is responsible for maintaining mitochondrial membrane potential in bloodstream form *Trypanosoma brucei*. Eukaryot Cell 5, 45–53, doi:10.1128/EC.5.1.45-53.2006 (2006).

41 van der Laan, M., Bechtluft, P., Kol, S., Nouwen, N. & Driessen, A. J. F1F0 ATP synthase subunit c is a substrate of the novel YidC pathway for membrane protein biogenesis. J Cell Biol 165, 213–222, doi:10.1083/jcb.200402100 (2004).

42 Jia, L., Dienhart, M. K. & Stuart, R. A. Oxa1 directly interacts with Atp9 and mediates its assembly into the mitochondrial F1Fo-ATP synthase complex. Mol Biol Cell 18, 1897–1908, doi:10.1091/mbc.e06-10-0925 (2007).

43 Hierro-Yap, C. et al. Bioenergetic consequences of F(o)F(1)-ATP synthase/ATPase deficiency in two life cycle stages of *Trypanosoma brucei*. J Biol Chem 296, 100357, doi:10.1016/j.jbc.2021.100357 (2021).

44 Pena-Diaz, P. et al. Trypanosomal mitochondrial intermediate peptidase does not behave as a classical mitochondrial processing peptidase. PLoS One 13, e0196474, doi:10.1371/journal.pone.0196474 (2018).

45 Harsman, A. et al. The non-canonical mitochondrial inner membrane presequence translocase of trypanosomatids contains two essential rhomboid-like proteins. Nat Commun 7, 13707, doi:10.1038/ncomms13707 (2016).

46 Hu, Y. et al. A Supercomplex Incorporating Both Electron Transport Chain and ATP Synthase. doi:10.65215/LTSpreprints.2026.04.03.000171 (2026).

47 Montgomery, M. G., Gahura, O., Leslie, A. G. W., Zikova, A. & Walker, J. E. ATP synthase from *Trypanosoma brucei* has an elaborated canonical F-1-domain and conventional catalytic sites. P Natl Acad Sci USA 115, 2102–2107, doi:10.1073/pnas.1720940115 (2018).

48 Zhou, D. et al. Protocol for mitochondrial isolation and sub-cellular localization assay for mitochondrial proteins. STAR Protoc 4, 102088, doi:10.1016/j.xpro.2023.102088 (2023).

49 Panigrahi, A. K. et al. A comprehensive analysis of *Trypanosoma brucei* mitochondrial proteome. Proteomics 9, 434–450, doi:10.1002/pmic.200800477 (2009).

50 Peikert, C. D. et al. Charting organellar importomes by quantitative mass spectrometry. Nat Commun 8, 15272, doi:10.1038/ncomms15272 (2017).

51 Niemann, M. et al. Mitochondrial outer membrane proteome of *Trypanosoma brucei* reveals novel factors required to maintain mitochondrial morphology. Mol Cell Proteomics 12, 515–528, doi:10.1074/mcp.M112.023093 (2013).

52 Zikova, A., Verner, Z., Nenarokova, A., Michels, P. A. M. & Lukes, J. A paradigm shift: The mitoproteomes of procyclic and bloodstream *Trypanosoma brucei* are comparably complex. PLoS Pathog 13, e1006679, doi:10.1371/journal.ppat.1006679 (2017).

53 von Kanel, C. et al. Homologue replacement in the import motor of the mitochondrial inner membrane of trypanosomes. Elife 9, doi:10.7554/eLife.52560 (2020).

54 Hildenbeutel, M. et al. The membrane insertase Oxa1 is required for efficient import of carrier proteins into mitochondria. J Mol Biol 423, 590–599, doi:10.1016/j.jmb.2012.07.018 (2012).

55 Jia, L. et al. Yeast Oxa1 interacts with mitochondrial ribosomes: the importance of the C-terminal region of Oxa1. EMBO J 22, 6438–6447, doi:10.1093/emboj/cdg624 (2003).

56 Ramrath, D. J. F. et al. Evolutionary shift toward protein-based architecture in trypanosomal mitochondrial ribosomes. Science 362, doi:10.1126/science.aau7735 (2018).

57 Skodova-Sverakova, I., Horvath, A. & Maslov, D. A. Identification of the mitochondrially encoded subunit 6 of F1FO ATPase in *Trypanosoma brucei*. Mol Biochem Parasitol 201, 135–138, doi:10.1016/j.molbiopara.2015.08.002 (2015).

58 Schneider, H. C., Westermann, B., Neupert, W. & Brunner, M. The nucleotide exchange factor MGE exerts a key function in the ATP-dependent cycle of mt-Hsp70-Tim44 interaction driving mitochondrial protein import. EMBO J 15, 5796–5803 (1996).

59 Schulz, C., Schendzielorz, A. & Rehling, P. Unlocking the presequence import pathway. Trends Cell Biol 25, 265–275, doi:10.1016/j.tcb.2014.12.001 (2015).

60 Esser, K., Tursun, B., Ingenhoven, M., Michaelis, G. & Pratje, E. A novel two-step mechanism for removal of a mitochondrial signal sequence involves the mAAA complex and the putative rhomboid protease Pcp1. J Mol Biol 323, 835–843, doi:10.1016/s0022-2836(02)01000-8 (2002).

61 McQuibban, G. A., Saurya, S. & Freeman, M. Mitochondrial membrane remodelling regulated by a conserved rhomboid protease. Nature 423, 537–541, doi:10.1038/nature01633 (2003).

62 Herlan, M., Vogel, F., Bornhovd, C., Neupert, W. & Reichert, A. S. Processing of Mgm1 by the rhomboid-type protease Pcp1 is required for maintenance of mitochondrial morphology and of mitochondrial DNA. J Biol Chem 278, 27781–27788, doi:10.1074/jbc.M211311200 (2003).

63 Hashimi, H., Gahura, O. & Panek, T. Bringing together but staying apart: decisive differences in animal and fungal mitochondrial inner membrane fusion. Biol Rev Camb Philos Soc 100, 920–935, doi:10.1111/brv.13168 (2025).

64 Amutha, B., Gordon, D. M., Gu, Y. & Pain, D. A novel role of Mgm1p, a dynamin-related GTPase, in ATP synthase assembly and cristae formation/maintenance. Biochem J 381, 19–23, doi:10.1042/BJ20040566 (2004).

65 Urban, S., Lee, J. R. & Freeman, M. Drosophila rhomboid-1 defines a family of putative intramembrane serine proteases. Cell 107, 173–182, doi:10.1016/s0092-8674(01)00525-6 (2001).

66 Lemberg, M. K. & Freeman, M. Functional and evolutionary implications of enhanced genomic analysis of rhomboid intramembrane proteases. Genome Res 17, 1634–1646, doi:10.1101/gr.6425307 (2007).

67 Kolli, R. et al. The OXA2a Insertase of *Arabidopsis* Is Required for Cytochrome c Maturation. Plant Physiol 184, 1042–1055, doi:10.1104/pp.19.01248 (2020).

68 Funes, S. et al. Independent gene duplications of the YidC/Oxa/Alb3 family enabled a specialized cotranslational function. Proc Natl Acad Sci U S A 106, 6656–6661, doi:10.1073/pnas.0809951106 (2009).

69 Gerdes, L. et al. A second thylakoid membrane-localized Alb3/OxaI/YidC homologue is involved in proper chloroplast biogenesis in *Arabidopsis thaliana*. J Biol Chem 281, 16632–16642, doi:10.1074/jbc.M513623200 (2006).

70 Ackermann, B. et al. Chloroplast Ribosomes Interact With the Insertase Alb3 in the Thylakoid Membrane. Front Plant Sci 12, 781857, doi:10.3389/fpls.2021.781857 (2021).

71 Kohler, A., Barrientos, A., Fontanesi, F. & Ott, M. The functional significance of mitochondrial respiratory chain supercomplexes. EMBO Rep 24, e57092, doi:10.15252/embr.202357092 (2023).

72 Muhleip, A. W., Dewar, C. E., Schnaufer, A., Kuhlbrandt, W. & Davies, K. M. In situ structure of trypanosomal ATP synthase dimer reveals a unique arrangement of catalytic subunits. Proc Natl Acad Sci U S A 114, 992–997, doi:10.1073/pnas.1612386114 (2017).

73 Wu, M. et al. How respiratory complexes and ATP synthase co-assemble to build cristae doi:10.65215/LTSpreprints.2026.04.03.000172 (2026).

74 Kohler, R. et al. YidC and Oxa1 form dimeric insertion pores on the translating ribosome. Mol Cell 34, 344–353, doi:10.1016/j.molcel.2009.04.019 (2009).

75 Englmeier, R., Pfeffer, S. & Forster, F. Structure of the Human Mitochondrial Ribosome Studied In Situ by Cryoelectron Tomography. Structure 25, 1574–1581 e1572, doi:10.1016/j.str.2017.07.011 (2017).

76 Pfeffer, S., Woellhaf, M. W., Herrmann, J. M. & Forster, F. Organization of the mitochondrial translation machinery studied in situ by cryoelectron tomography. Nat Commun 6, 6019, doi:10.1038/ncomms7019 (2015).

77 Wang, S. et al. Structural basis of TACO1-mediated efficient mitochondrial translation. Nat Commun 17, doi:10.1038/s41467-026-69156-y (2026).

78 Hartl, F. U., Schmidt, B., Wachter, E., Weiss, H. & Neupert, W. Transport into mitochondria and intramitochondrial sorting of the Fe/S protein of ubiquinol-cytochrome c reductase. Cell 47, 939–951, doi:10.1016/0092-8674(86)90809-3 (1986).

79 Hell, K., Herrmann, J., Pratje, E., Neupert, W. & Stuart, R. A. Oxa1p mediates the export of the N- and C-termini of pCoxII from the mitochondrial matrix to the intermembrane space. FEBS Lett 418, 367–370, doi:10.1016/s0014-5793(97)01412-9 (1997).

80 Gahura, O. & Chauhan, P. Mitochondrial ribosomes in apicomplexan and trypanosomatid parasites: Dissimilar drivers of complexity and convergent features. PLoS Pathog 22, e1013920, doi:10.1371/journal.ppat.1013920 (2026).

81 Gahura, O., Chauhan, P. & Zikova, A. Mechanisms and players of mitoribosomal biogenesis revealed in trypanosomatids. Trends Parasitol 38, 1053–1067, doi:10.1016/j.pt.2022.08.010 (2022).

82 Hashimi, H. A parasite’s take on the evolutionary cell biology of MICOS. PLoS Pathog 15, e1008166, doi:10.1371/journal.ppat.1008166 (2019).

83 Wong, J. E., Zikova, A. & Gahura, O. The Ancestral Shape of the Access Proton Path of Mitochondrial ATP Synthases Revealed by a Split Subunit-a. Mol Biol Evol 40, doi:10.1093/molbev/msad146 (2023).

84 Altschul, S. F., Gish, W., Miller, W., Myers, E. W. & Lipman, D. J. Basic local alignment search tool. J Mol Biol 215, 403–410, doi:10.1016/S0022-2836(05)80360-2 (1990).

85 van Kempen, M. et al. Fast and accurate protein structure search with Foldseek. Nat Biotechnol, doi:10.1038/s41587-023-01773-0 (2023).

86 Abramson, J. et al. Accurate structure prediction of biomolecular interactions with AlphaFold 3. Nature 630, 493–500, doi:10.1038/s41586-024-07487-w (2024).

87 Hallgren, J. et al., doi:10.1101/2022.04.08.487609 (2022).

88 Meng, E. C. et al. UCSF ChimeraX: Tools for structure building and analysis. Protein Sci 32, e4792, doi:10.1002/pro.4792 (2023).

89 Richter, D. J. et al. EukProt: A database of genome-scale predicted proteins across the diversity of eukaryotes. Peer Community Journal 2, doi:10.24072/pcjournal.173 (2022).

90 Frickey, T. & Lupas, A. CLANS: a Java application for visualizing protein families based on pairwise similarity. Bioinformatics 20, 3702–3704, doi:10.1093/bioinformatics/bth444 (2004).

91 Kuraku, S., Zmasek, C. M., Nishimura, O. & Katoh, K. aLeaves facilitates on-demand exploration of metazoan gene family trees on MAFFT sequence alignment server with enhanced interactivity. Nucleic Acids Res 41, W22–28, doi:10.1093/nar/gkt389 (2013).

92 Mistry, J., Finn, R. D., Eddy, S. R., Bateman, A. & Punta, M. Challenges in homology search: HMMER3 and convergent evolution of coiled-coil regions. Nucleic Acids Res 41, e121, doi:10.1093/nar/gkt263 (2013).

93 Waterhouse, A. et al. SWISS-MODEL: homology modelling of protein structures and complexes. Nucleic Acids Res 46, W296–W303, doi:10.1093/nar/gky427 (2018).

94 Criscuolo, A. & Gribaldo, S. BMGE (Block Mapping and Gathering with Entropy): a new software for selection of phylogenetic informative regions from multiple sequence alignments. BMC Evol Biol 10, 210, doi:10.1186/1471-2148-10-210 (2010).

95 Minh, B. Q. et al. IQ-TREE 2: New Models and Efficient Methods for Phylogenetic Inference in the Genomic Era. Mol Biol Evol 37, 1530–1534, doi:10.1093/molbev/msaa015 (2020).

96 Hoang, D. T., Chernomor, O., von Haeseler, A., Minh, B. Q. & Vinh, L. S. UFBoot2: Improving the Ultrafast Bootstrap Approximation. Mol Biol Evol 35, 518–522, doi:10.1093/molbev/msx281 (2018).

97 Guindon, S. et al. New algorithms and methods to estimate maximum-likelihood phylogenies: assessing the performance of PhyML 3.0. Syst Biol 59, 307–321, doi:10.1093/sysbio/syq010 (2010).

98 Alves, A. A. et al. Control of assembly of extra-axonemal structures: the paraflagellar rod of trypanosomes. J Cell Sci 133, doi:10.1242/jcs.242271 (2020).

99 Beneke, T. & Gluenz, E. LeishGEdit: A Method for Rapid Gene Knockout and Tagging Using CRISPR-Cas9. Methods Mol Biol 1971, 189–210, doi:10.1007/978-1-4939-9210-2_9 (2019).

100 Johnston, H. E. et al. Solvent Precipitation SP3 (SP4) Enhances Recovery for Proteomics Sample Preparation without Magnetic Beads. Anal Chem 94, 10320–10328, doi:10.1021/acs.analchem.1c04200 (2022).

101 Zikova, A., Husova, M., Sever, A., Kunzova, M. & Dolezelova, E. Assessment of Mitochondrial Membrane Potential in Intact and Detergent-Permeabilized *Trypanosoma brucei* Insect and Bloodstream Forms. Methods Mol Biol 3014, 325–347, doi:10.1007/978-1-0716-5146-9_21 (2026).

102 Horvath, A., Nebohacova, M., Lukes, J. & Maslov, D. A. Unusual polypeptide synthesis in the kinetoplast-mitochondria from *Leishmania tarentolae*. Identification of individual de novo translation products. J Biol Chem 277, 7222–7230, doi:10.1074/jbc.M109715200 (2002).

