## Supplementary_materials for "Distinct roles of three trypanosomal Oxa1 insertases in biogenesis of mitochondrial membrane complexes"

**List of Supplementary Information:**

**Supplementary Table 1.** Mass spectrometry proteomic data from submitochondrial fractionation *(MS Excel file)*

**Supplementary Table 2.** Mass spectrometry proteomic data from immunoprecipitation experiments *(MS Excel file)*

**Supplementary Table 3.** List of oligonucleotides and antibodies *(MS Excel file)*

**Supplementary Figure 1.** Sequence alignment of TbOxa1 proteins and selected homologs.

**Supplementary Figure 2.** Evolutionary relationship of Oxa1 proteins in euglenozoans and selected homologs in other organisms.

**Supplementary Figure 3.** Characterization of cell lines with V5-tagged TbOxa1 paralogs.

**Supplementary Figure 4.** Verification of TbOxa1 bloodstream form knock out cell lines and phenotypes of double TbOxa1 knock outs.

**Supplementary Figure 5.** Verification of TbOxa1 procyclic knock out cell lines.

**Supplementary Figure 6.** Effect of TbOxa1-3 ablation on mitochondrial proteome.

**Supplementary Figure 7.** Identification of paralogs of ATP synthase subunit g and characterization of TbRPLP^KO^ strain.

**SUPPLEMENTARY FIGURES**

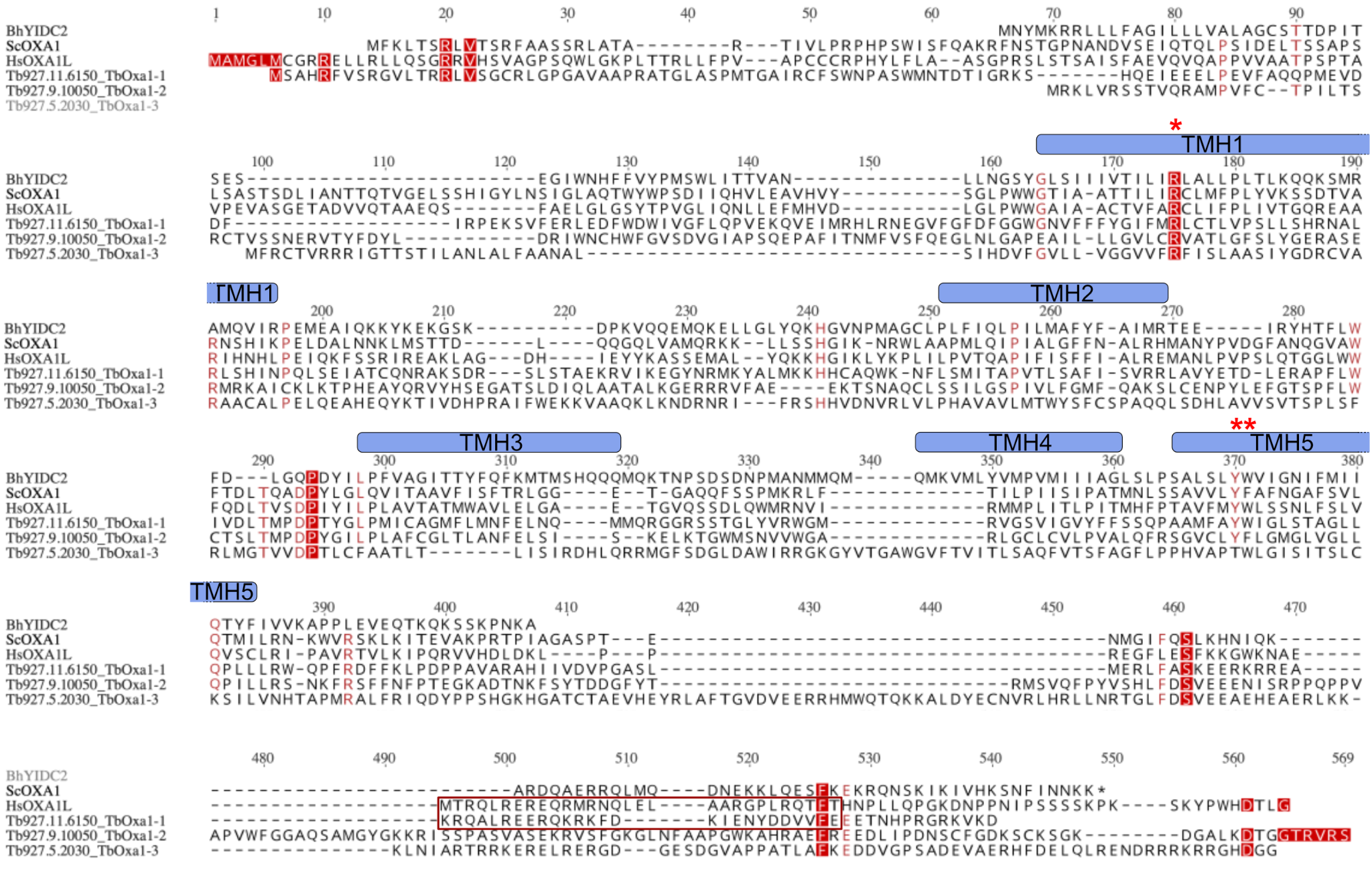

**Supplementary Figure 1. Sequence alignment of TbOxa1 proteins and selected homologs.** Multiple sequence alignment of the three TbOxa1 proteins with homologous proteins in *Bacillus halodurans* (BhYidC2), *Saccharomyces cerevisiae* (ScOxa1), and *Homo sapiens* (HsOXA1L). The transmembrane helices (TMH) 1 to 5 of BhYidC2^1^ are indicated above the sequences. The residues essential for insertase activity in BhYidC2 are marked with red asterisks. The region corresponding to the mitoribosome binding site 2 in HsOXA1L^2^ and corresponding region in TbOxa1-1 is indicated in a red box.

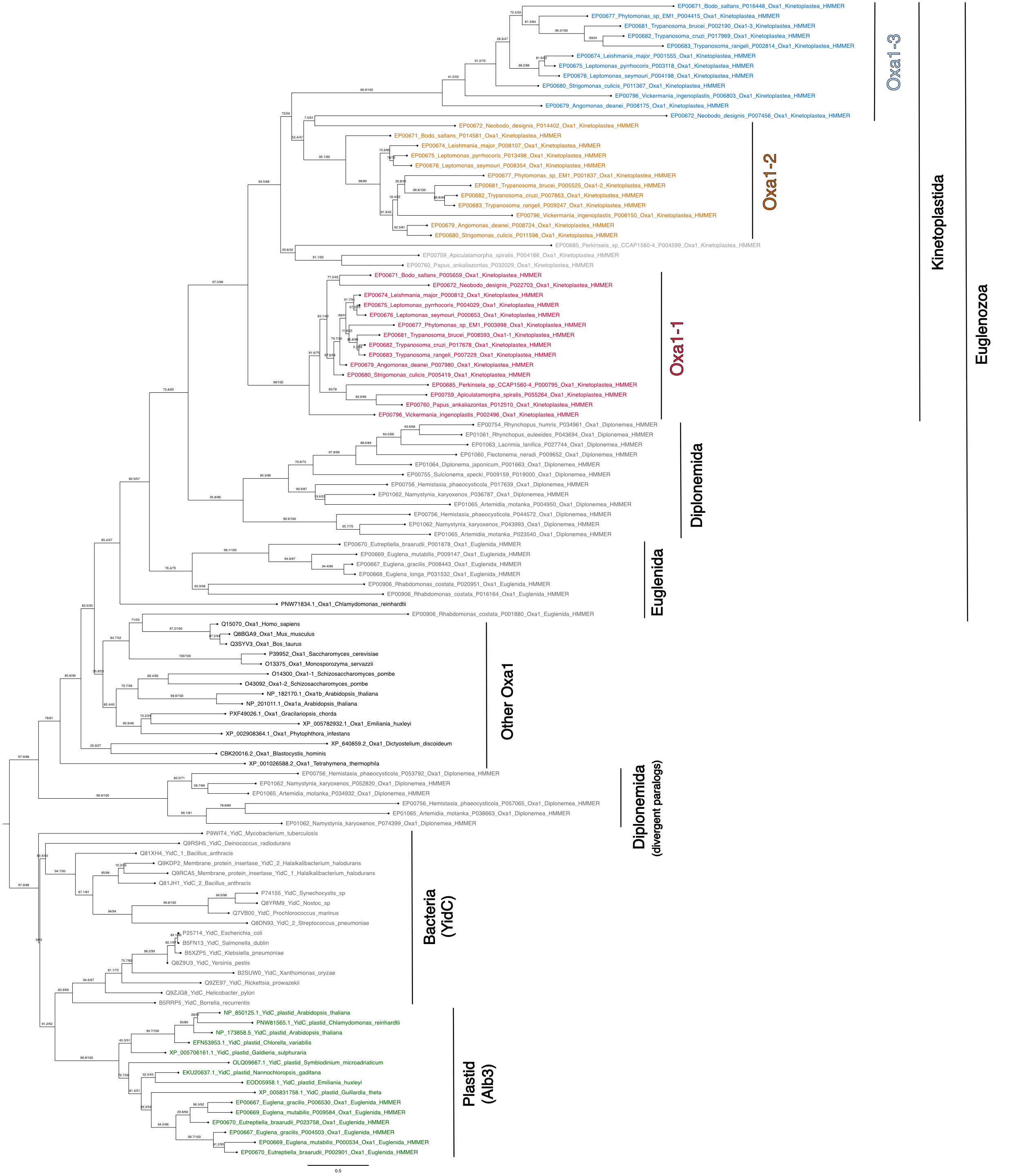

**Supplementary Figure 2. Evolutionary relationship of Oxa1 proteins in euglenozoans and selected homologs in other organisms.** The maximum likelihood phylogenetic tree was rooted between bacterial YidC and eukaryotic Oxa1 clusters.

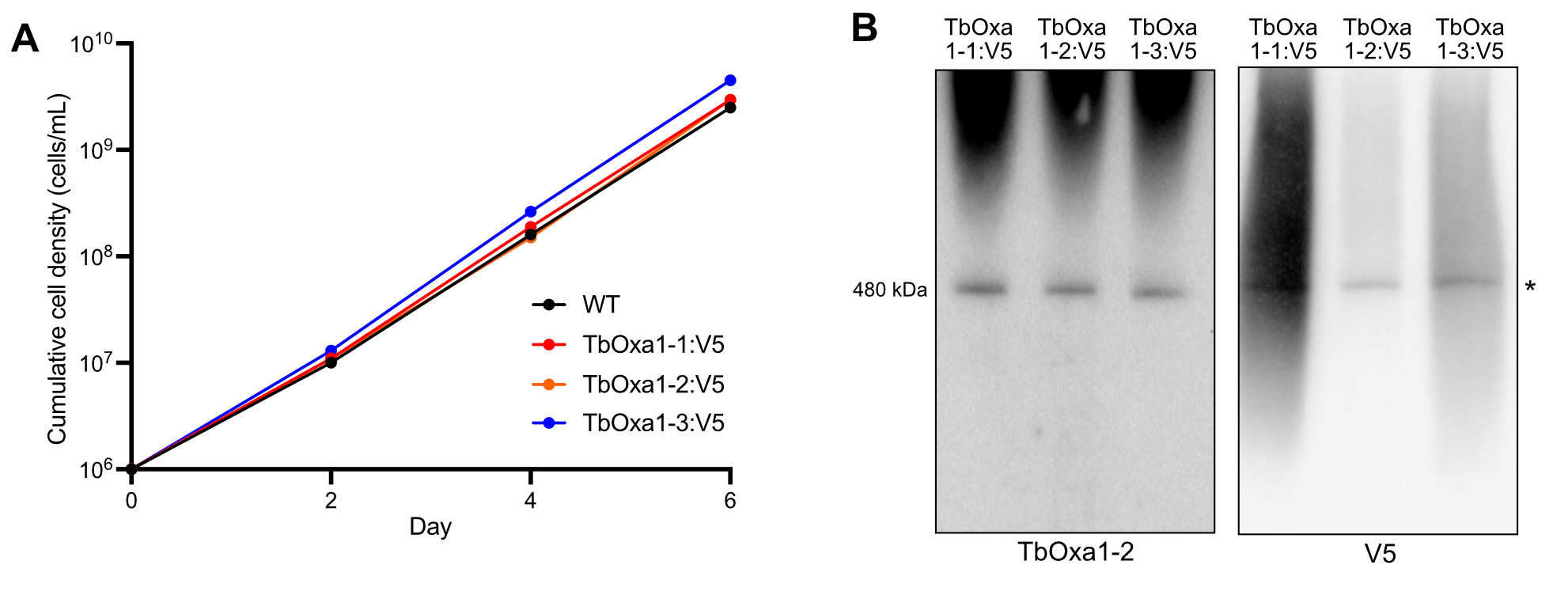

**Supplementary Figure 3. Characterization of cell lines with V5-tagged TbOxa1 paralogs. (A)** Growth curves of WT and V5-tagged TbOxa1-expressing *T. brucei* (TbOxa1:V5) PF cell lines grown in SDM-79 medium. **(B)** Immunoblot of mitochondrial lysates from WT and TbOxa1:V5 PF cells resolved by BN-PAGE probed with antibodies against TbOxa1-2 and the V5 epitope.

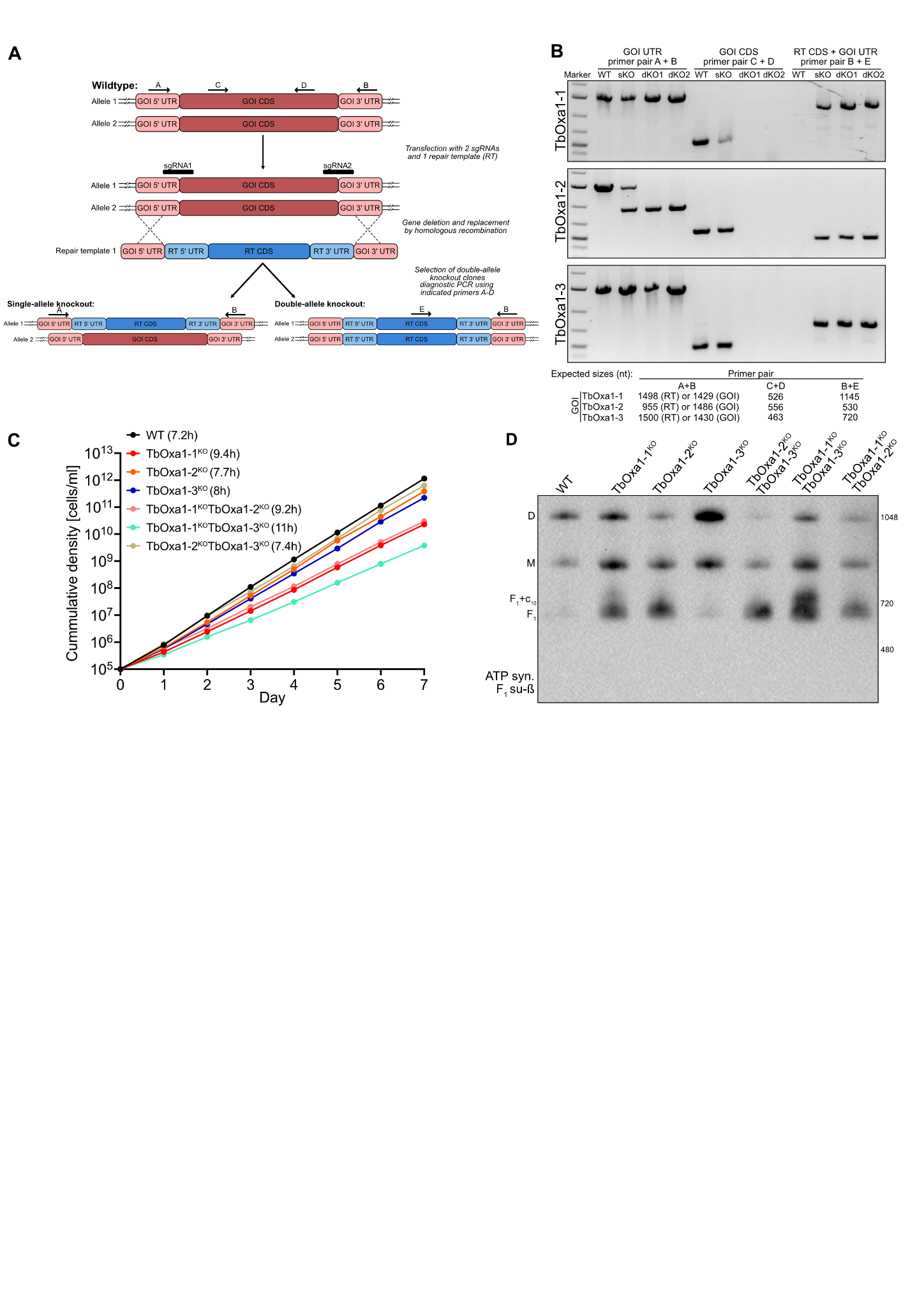

**Supplementary Figure 4. Verification of TbOxa1 bloodstream form knock out cell lines and phenotypes of double TbOxa1 knock outs. (A)** A scheme of knocking out the TbOxa1 genes in Cas9-expressing *T. brucei* BF cells by CRISPR/Cas9 strategy. The single guide RNA molecules (sgRNA1 and 2) targeting the regions flanking the coding sequence of genes of interest (GOI CDS) are indicated. The positions of primers A to E used for verification of the deletion of GOI CDS and its replacement by an antibiotic resistance conferring gene coding sequence repair template (RT CDS) are indicated. **(B)** Agarose gel electrophoresis of PCR products using indicated primer pairs to verify TbOxa1 gene deletion and replacement with RT CDS. Parental strains (WT) were used as a negative control. Expected sizes of PCR products for each primer pair combination are indicated below. sKO - single allele knock-out; dKO - both alleles knock-out. **(C)** Growth curve of WT and indicated single or double gene TbOxa1 knockout BF cell lines. **(D)** Immunoblot of mitochondrial lysates from WT and indicated single or double gene BF cells resolved by BN-PAGE probed with antibody against ATP synthase subunit β.

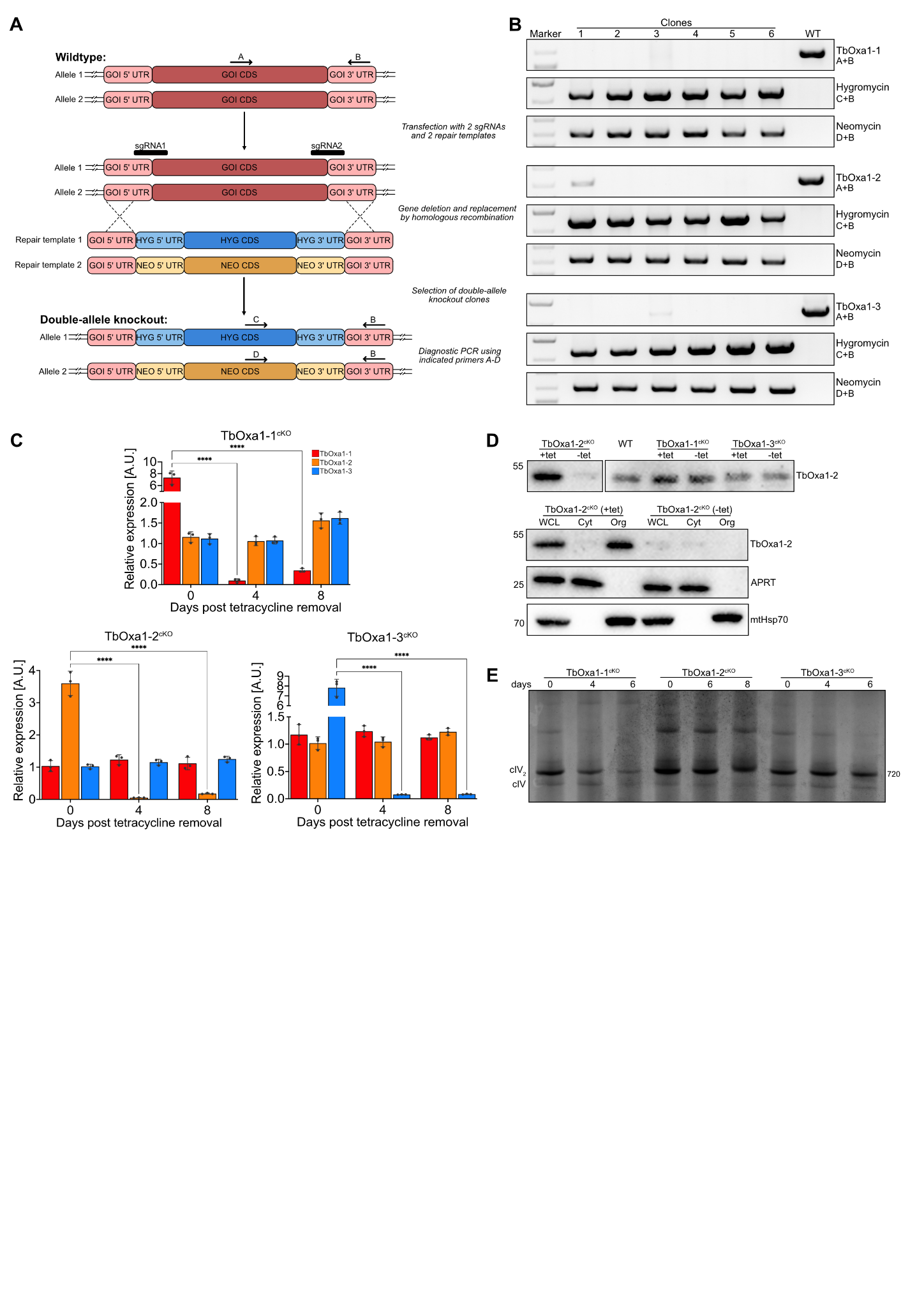

**Supplementary Figure 5. Verification of TbOxa1 procyclic knock out cell lines. (A)** A scheme of knocking out the TbOxa1 genes in Cas9-expressing *T. brucei* procyclic cells by CRISPR/Cas9 strategy. The single guide RNA molecules (sgRNA1 and 2) targeting the regions flanking the coding region of our genes of interest (GOI CDS) are indicated. The positions of primers A to D used for verification of the deletion of GOI CDS and its replacement by hygromycin or neomycin resistance conferring gene coding sequence repair template (HYG/NEO CDS) are indicated. **(B)** Agarose gel electrophoresis of PCR products using indicated primer pairs to verify TbOxa1 gene deletion and replacement with RT CDS (hygromycin and neomycin) in WT and select PF clones. **(C)** Quantification of TbOxa1 transcript levels by RT-qPCR in TbOxa1^cKO^ procyclic cells at indicated timepoints after tetracycline removal. Values were normalized to expression of reference gene tubulin and to expression of the respective TbOxa1 paralog in wild-type cells. **** indicates p-value <0.0001. **(D)** Upper panel shows immunoblots of whole cell lysates from WT and TbOxa1^cKO^ procyclic cell lines in presence (+tet) of tetracycline or 4 days after tetracycline removal (-tet) resolved by SDS-PAGE and probed with antibody against TbOxa1-2. The lower panel shows immunoblots of lysates from whole cells (WCL) and cytosolic (Cyt) and organellar (Org) fractions from TbOxa1-2^cKO^ cells in presence and absence of tetracycline. **(E)** BN-PAGE gel with resolved mitochondrial lysates from TbOxa1^cKO^ procyclic cell lines from indicated days after tetracycline removal stained for complex IV activity.

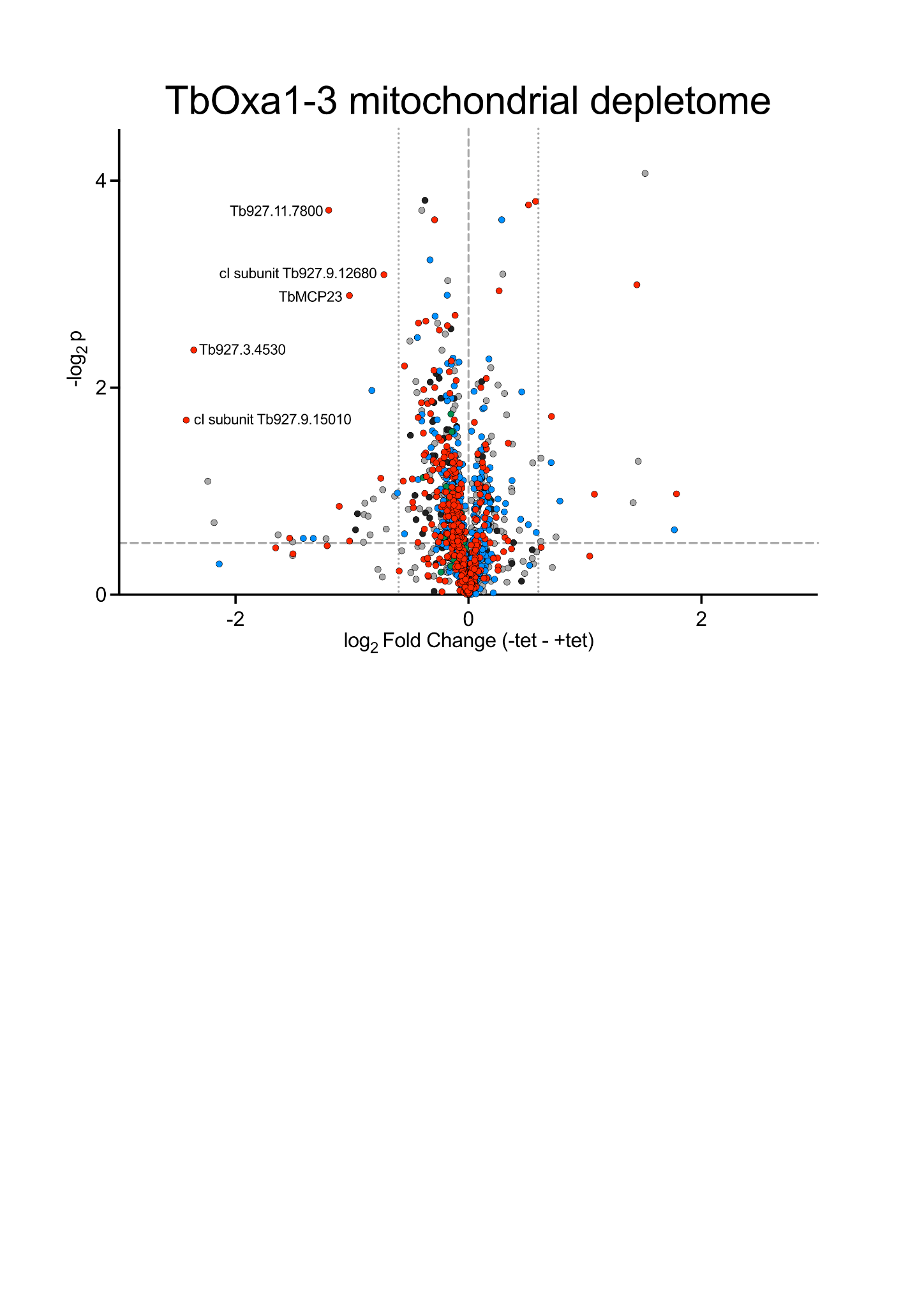

**Supplementary Figure 6. Effect of TbOxa1-3 ablation on mitochondrial proteome.** Dot plot showing the changes in protein content in the mitochondria from TbOxa1-3^cKO^ procyclic cells expressing the ectopic copy (+tet) and the same cells 6 days after tetracycline removal (-tet).

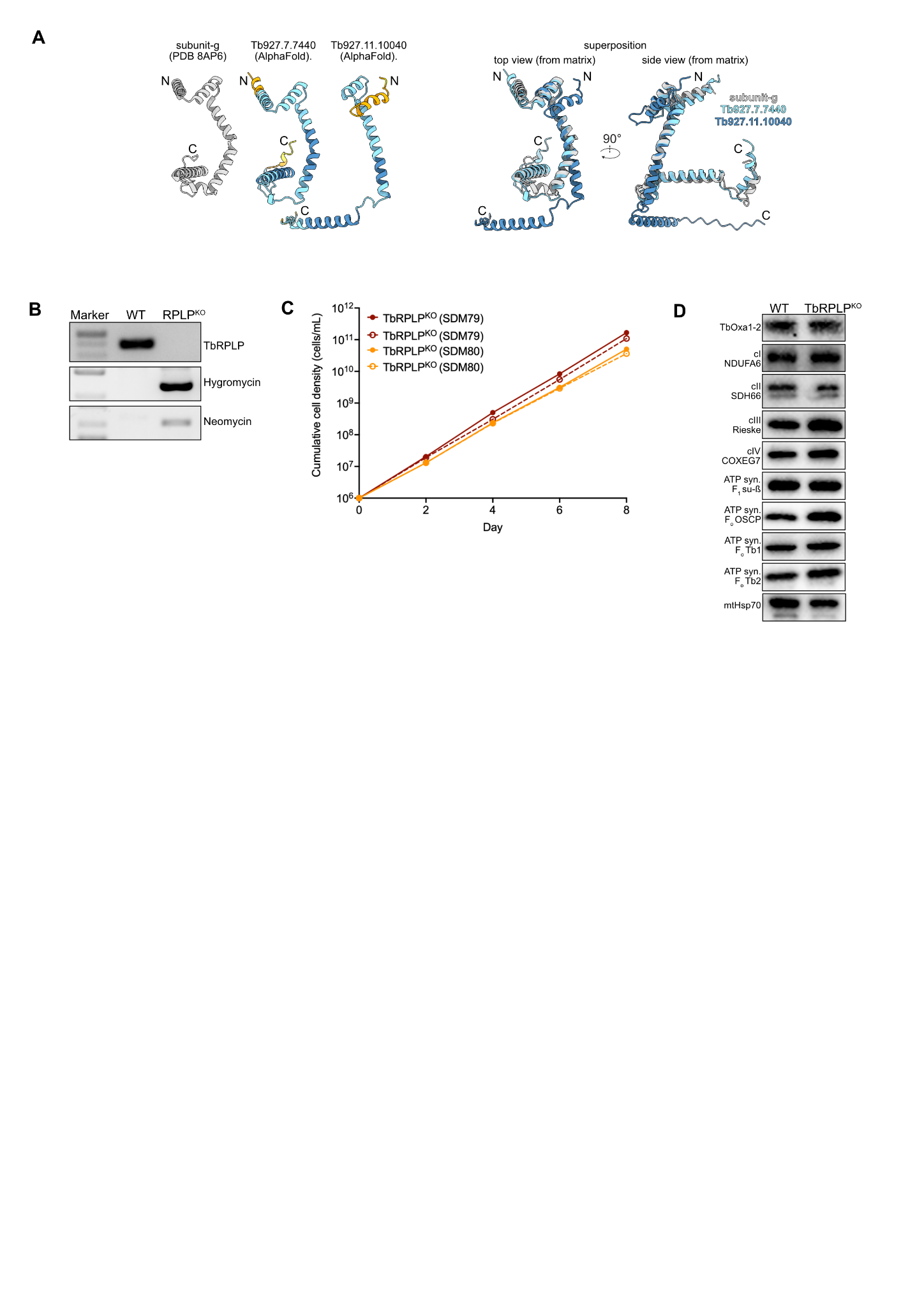

**Supplementary Figure 7. Identification of paralogs of ATP synthase subunit g and characterization of TbRPLP^KO^ strain. (A)** Comparison of AlphaFold predicted structures of identified paralogs of ATP subunit g (TriTrypDB accession numbers Tb927.7.7440 and Tb927.11.10040) with the structure of canonical subunit g as resolved by cryoEM in complete dimer of ATP synthase. In the left panel, subunit g paralogs are colored by pLDDT using the standard AlphaFold palette. **(B)** Verification of TbRPLP knock out by PCR with primer pairs detecting indicated coding sequences. **(C)** Growth curve of TbRPLP^KO^ cells in SMD79 and SDM80 media. **(D)** Steady state levels of indicated proteins in TbRPLP^KO^ strain assayed by immunoblotting of mitochondrial lysates resolved by SDS-PAGE.

**Supplementary References**

1	Kumazaki, K. *et al.* Structural basis of Sec-independent membrane protein insertion by YidC. *Nature* **509**, 516-520, doi:10.1038/nature13167 (2014).

2	Itoh, Y. *et al.* Mechanism of membrane-tethered mitochondrial protein synthesis. *Science* **371**, 846-849, doi:10.1126/science.abe0763 (2021).
